# Joint single cell DNA-Seq and RNA-Seq of cancer reveals subclonal signatures of genomic instability and gene expression

**DOI:** 10.1101/445932

**Authors:** Noemi Andor, Billy T. Lau, Claudia Catalanotti, Vijay Kumar, Anuja Sathe, Kamila Belhocine, Tobias D. Wheeler, Andrew D. Price, Maengseok Song, Željko Džakula, Jon Sorenson, David Stafford, Zachary Bent, Laura DeMare, Lance Hepler, Susana Jett, Bill Kengli Lin, Shamoni Maheshwari, Anthony J. Makarewicz, Mohammad Rahimi, Sanjam S. Sawhney, Martin Sauzade, Joe Shuga, Katrina Sullivan-Bibee, Adam Weinstein, Wei Yang, Yifeng Yin, Matthew Kubit, Jiamin Chen, Susan M. Grimes, Carlos Jose Suarez, George A. Poultsides, Michael Schnall-Levin, Rajiv Bharadwaj, Hanlee P. Ji

## Abstract

Sequencing the genomes of individual cancer cells provides the highest resolution of intratumoral heterogeneity. To enable high throughput single cell DNA-Seq across thousands of individual cells per sample, we developed a droplet-based, automated partitioning technology for whole genome sequencing. We applied this approach on a set of gastric cancer cell lines and a primary gastric tumor. In parallel, we conducted a separate single cell RNA-Seq analysis on these same cancers and used copy number to compare results. This joint study, covering thousands of single cell genomes and transcriptomes, revealed extensive cellular diversity based on distinct copy number changes, numerous subclonal populations and in the case of the primary tumor, subclonal gene expression signatures. We found genomic evidence of positive selection – where the percentage of replicating cells per clone is higher than expected – indicating ongoing tumor evolution. Our study demonstrates that joining single cell genomic DNA and transcriptomic features provides novel insights into cancer heterogeneity and biology.

**SIGNIFICANCE:** We conducted a massively parallel DNA sequencing analysis on a set of gastric cancer cell lines and a primary gastric tumor in combination with a joint single cell RNA-Seq analysis. This joint study, covering thousands of single cell genomes and transcriptomes, revealed extensive cellular diversity based on distinct copy number changes, numerous subclonal populations and in the case of the primary tumor, subclonal gene expression signatures. We found genomic evidence of positive selection where the percentage of replicating cells per clone is higher than expected indicating ongoing tumor evolution. Our study demonstrates that combining single cell genomic DNA and transcriptomic features provides novel insights into cancer heterogeneity and biology.

---

Single cell DNA sequencing **(scDNA-Seq)** identifies somatic genetic alterations such as somatic copy number variants **(CNVs)**. For cancer, single cell CNVs paint a high-resolution profile of intratumoral heterogeneity and subclonal structure (1–4) present in primary tumors (5, 6), metastases (7, 8), patient-derived xenografts and even cancer cell lines (9). This underlying genomic variation seen among a cancer’s subclonal populations provides a “fuel” for tumor evolution and adaptation to ongoing therapy. Notably, the dominant subclones of resistant tumors (5, 6), metastases (7, 8), patient-derived xenografts and cell lines (9) often originate from minor subclones in the primary tumor.

The prevalence of intratumoral heterogeneity has implications for cancer biology studies. Cancer cell lines are used to model tumor growth, evaluate metastatic potential and determine drug sensitivities. However, cancer cell lines have subpopulations with extensive fitness diversity (9, 10). This may lead to different drug responses within the same cell line (10). The application of single cell DNA-Seq quantifies the extent of subclonal diversity and may impact these type of studies.

This scDNA-Seq approach relies on either low coverage whole genome sequencing **(WGS)** to identify somatic CNVs or targeted sequencing to identify cancer mutations (1–4, 11, 12). However, the cellular throughput of scDNA-Seq has been limited, with a typical maximum of hundred cells. Greater sampling of tumor tissues provides an opportunity to expand the scope of intratumoral characterization. However, increasing cellular sequencing throughput is difficult for a number of reasons: limitations of single cell partitioning methods whether plate-based or using flow cytometry isolation (4); issues with amplifying genomic DNA from single cells; complex methods for isolating nuclei; intricate enzymology steps for library preparation (13, 14). As a solution that enables massive scale scDNA-Seq, we developed a droplet-based partitioning technology that rapidly processes thousands of cells per sample for library preparation in a highly automated fashion. Using this new approach, we conducted single cell WGS on thousands of cells for nine gastric cancer cell lines and a primary gastric tumor. This extensive cellular sampling provided robust characterization of subclonal structure of gastric cancer, determined cell cycle assignments and identified quantitative features related to tumor cell selection.

The majority of single cell genomic studies of cancer have focused on the use of single cell RNA-Seq **(scRNA-Seq)**, where one sequences thousands of individual transcriptomes from a given tumor (3, 15–17). By conducting a joint scDNA-Seq and scRNA-Seq analysis, one identifies underlying genomic alterations among the cells in the sample, subclonal cellular diversity and transcriptome features indicative of differences in biological pathways among the cellular populations. There are only a few studies, such as published by Kim et al., which combine both single cell methods for studying cancer (3, 15–18). Technical challenges limit the number of cells for WGS analysis and conducting joint studies, outside of cancer, often rely on specific large cell types, such as oocytes that are readily manipulated (15).

To conduct a parallel, joint scDNA-Seq and scRNA-Seq study of cancer cells, we also performed a separate, large-scale single cell transcriptome analysis of the same ten cancers **(Fig. 1*A*)**. Analyzing over 30,000 cellular transcriptomes form this set of cancers, we determined CNVs based on scRNA-Seq. Subsequently, we compared the CNV-defined subclonal populations detected from both scRNA-Seq and scDNA-Seq and in the case of the primary gastric tumor, overlaid the data to ascribe transcriptional features to subclonal populations.

**Figure 1.**
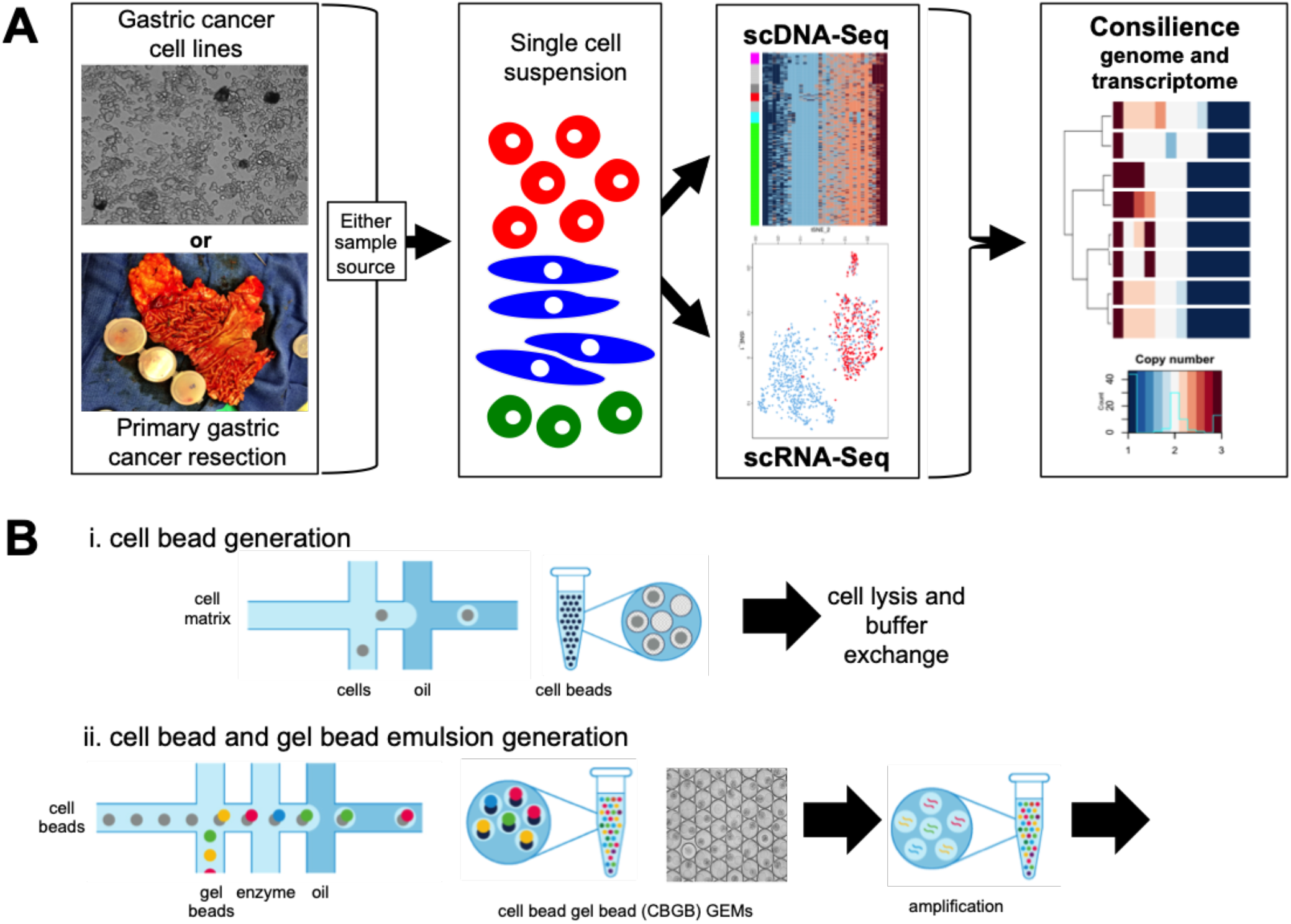
Single cell sequencing strategy. **(A)** Study design. Aliquots of cell suspension were used to conduct separate scRNA-Seq and scDNA-Seq analysis. CNVs are called independently from scDNA- and scRNA-Seq results and used to identify and mutually validate coexisting clones within each sample. ScRNA-Seq informs what genes each clone expresses, while scDNA-Seq has a higher resolution on the genomic instability of each clone. **(B)** Single cell DNA sequencing technology overview. (**i**) Cell beads (CBs) are generated by injection of cells into a microfluidic chip with a polymer matrix. Sub-nanoliter droplets are formed, with droplets containing either zero or one cell. After removal of droplets from the microfluidic cartridge, the cell beads form a crosslinked hydrogel bead. After emulsion breaking, the hydrogel CBs remain intact. Lysis and buffer exchange are performed without loss of cellular genomic material. (ii) CBs are loaded alongside barcode-containing gel beads (GBs), which enables the identification of sequence data to the originating droplet partition. Sub-nanoliter droplets are generated containing one of each type of bead. Whole genome amplification and barcoding subsequently takes place, which results in barcode-tagged amplified genomic DNA. The emulsion is then broken and standard library preparation procedures are performed to generate an Illumina sequencing library.

## RESULTS

### High Throughput Processing for Single Cell DNA-Seq

For the isolation of large numbers of single cells during library preparation, we developed a two-stage microfluidic droplet-based technology for the automatic generation of high cell number scDNA-Seq libraries. Similar to linked-read sequencing for genome phasing (19) and single cell transcriptome analysis (20), microfluidic droplets were loaded with a barcoded hydrogel bead that tags DNA. This feature enabled the tracking of sequence reads originating from individual droplets and their analyte molecules. In a first microfluidic chip, individual cells were encapsulated with paramagnetic particles and hydrogel matrix precursors to form cell-containing magnetic hydrogel beads, or ‘cell beads’ **(CBs)**. As in previous studies that load single cells into microfluidic droplets for transcriptomic studies (20), cellular suspensions were first loaded at Poisson limit dilution into droplets using a microfluidic chip. The resultant CBs contained either zero, one or multiple cells **(Fig. 1B)**.

After breaking the gelled CB emulsion, we used the magnetic properties of the encapsulating cell matrix to efficiently integrate microfluidic and macrofluidic processes to enable nuclear DNA processing for downstream amplification and barcoding. The CB hydrogel structure remains intact after emulsion breaking. The pore structure of the CB matrix facilitated the confinement of large genomic DNA molecules while keeping them diffusively accessible to lysis and denaturation agents. We used this CB feature to lyse cells, digest proteins, and denature DNA. This process yielded freely accessible DNA trapped in CBs suitable for re-partitioning in a second microfluidic step.

In a second microfluidic chip, processed CBs were injected into another microfluidic droplet generator cartridge for barcoding and single-cell whole genome amplification. A novel microfluidic system was employed whereby a single CB is encapsulated alongside a single barcoded gel bead **(GB)** with high efficiency leading to a cell bead-gel bead **(CBGBs)** emulsion **(Fig. 1*B*)**. The barcoded GB was similar to those used in previous studies for phasing genomes (19) and single cell transcriptome profiling (20) – an individual GB was functionalized with millions of copies of identical barcoded oligonucleotides that uniquely identifies a droplet during the sequencing reaction. The total number of unique droplet barcodes was approximately 737,000. The CBGB emulsion contained droplets with one, two, or three co-encapsulated beads. Up to 80% of droplets contained two beads consisting of only one CB and only one GB. The remainder consisted of a mixture of doubly loaded barcode GBs, doubly loaded CBs, or droplets loaded with three beads **(Fig. 1*B*)**.

After encapsulation and incubation, the CBGB emulsion was broken and underwent a modified library preparation protocol for Illumina sequencing **(Methods)**. Overall, the throughput of this microfluidicbased cellular isolation system demonstrated a scale up to tens of thousands of cells per a microfluidic chip. This processing capacity exceeded flow cytometry-based isolation by several orders of magnitude (20). Afterwards, single cell DNA libraries were sequenced with an Illumina system.

### Sequencing Genome Stable Diploid Cells

For a baseline, we evaluated genome stable diploid cells using peripheral blood mononuclear cells **(PBMCs)**. These studies enabled us to determine the extent of amplification bias, genomic dropout, and baseline ploidy calling performance. Data processing is described in the **Methods** section. We filtered aligned sequence reads specific to a single cell. The first 16 base pairs of Read 1 consist of a droplet-identifying barcode sequence. When counting the empirically observed distribution of reads per cell bead barcodes, we observed a strong bimodal distribution where there was a strong enrichment of reads belonging to less than 1 % of all observed barcode sequences in the dataset **(Fig. 2*A***). To identify droplet barcodes containing a single cell, we filtered outlier read counts and set a reads-per-barcode cutoff extrapolated form the barcode read maxima **(Methods, Fig. 2*A*)**. Approximately 96% of barcoded reads were assignable to a single cell, indicating the minimal spurious DNA contamination or amplification in other droplets.

**Figure 2.**
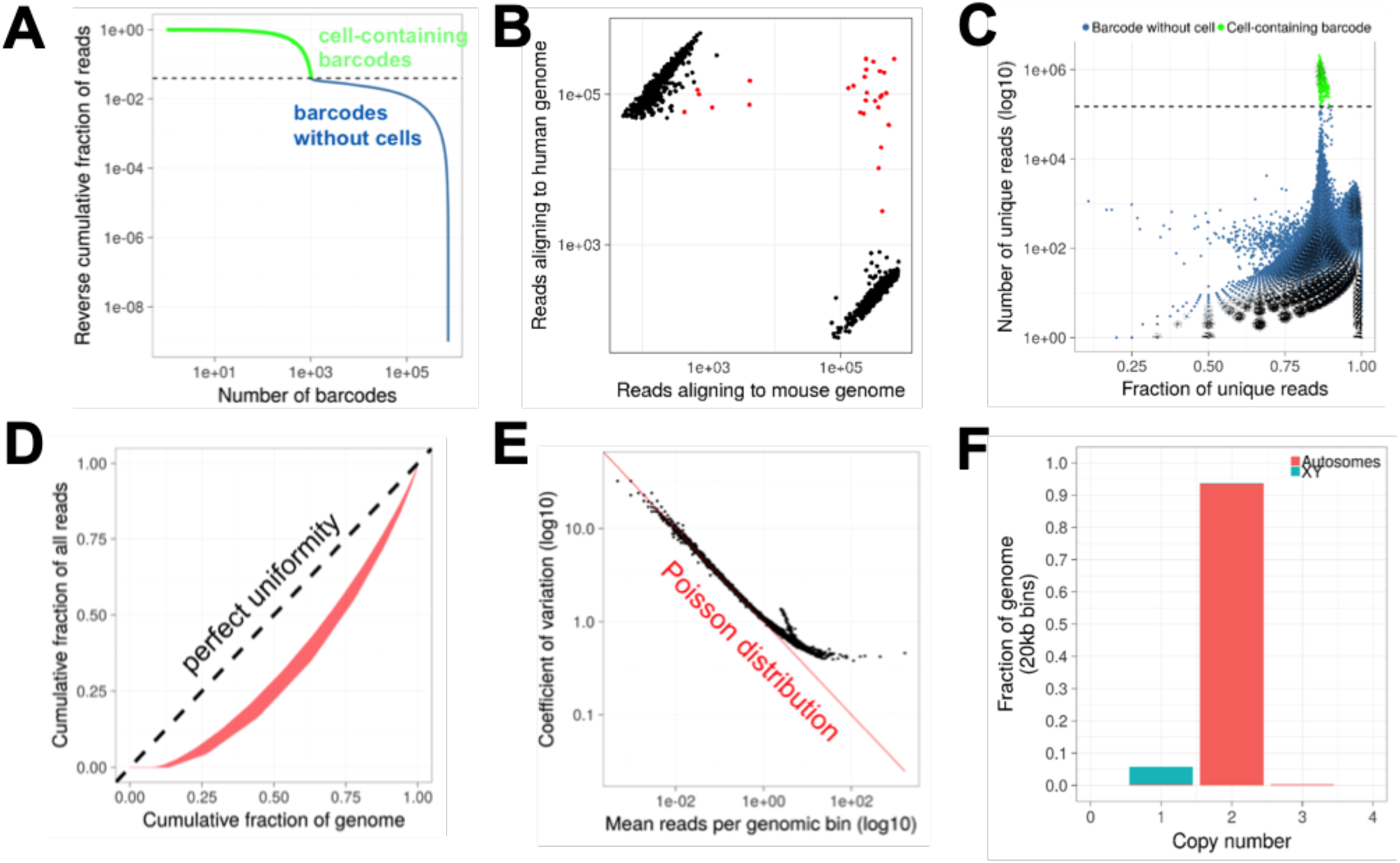
Technical performance of single cell DNA sequencing. **(A)** Sequence read distribution across droplet barcodes. Barcodes are assigned to be containing a cell based on measuring the quantiles of reads per observed barcode. Cell-containing barcodes are marked as a green line, and make up 96% of all sequence data. **(B)** scDNA-Seq of cell line mixtures. A mixture of human and mouse cell lines was used to generate a scDNA-Seq library. Each point represents a droplet partition. Black: droplet barcodes determined to be belonging to either only human or mouse genomes. Red: droplet barcodes with reads aligning to greater than 1% of both human and mouse genomes. **(C)** Unique reads in cell-containing barcodes. A scatterplot of the number of unique reads per barcode versus the fraction of unique reads per barcode is shown. Green points denote barcodes identified as those containing a cell. Dashed line: the cutoff read count for a barcode to be classified as being associated with a cell. **(D)** Uniformity of scDNA-Seq data. Lorenz curves of 20 PBMC cells selected at random. Curves significantly deviating from the diagonal reflect non-uniformity of amplification. **(E)** Reproducibility of scDNA-Seq across cells. Each point represents a genomic bin. The coefficient of variation of each genomic bin is plotted against its mean coverage across cells. A high dispersion would indicate significant reaction heterogeneity. **(F)** Histogram of copy number calls across 20kb bins in GRCh38. Copy number calls in 1,046 PBMCs were grouped according to either the autosomal or sex chromosomal regions.

To evaluate the performance of single cell loading per cell barcode, we analyzed a cellular mixture containing a mixture of human HEK-293T and mouse NIH-3T3 cells. After scDNA-Seq, doublet cells were identified by cellular barcodes with reads aligning to both mouse and human genomes. From a total of 1,313 cells, we observed that less than 2% of cells contained reads belonging to both species **(Fig. 2*B***), indicating that the vast majority of cells were loaded as single cells into CB emulsions. This result also indicated our ability to separate distinct cellular genomes.

Overall, we sequenced 1,046 diploid PBMCs with 1.2 billion 2×100 paired-end read pairs. The library had a median read duplication ratio of ~13% per cell, indicating a high overall complexity **(Fig. 2*C*)**. At least 100,000 reads per cell were sequenced, leading to at least one read per every 20kb genomic window **(*Fig. S*1 *A***). Generating Lorenz curves between the number of reads and fraction of genome sequenced, we determined that coverage demonstrated uniform amplification comparable to another method **(Fig. 2*D*)** (21). Finally, we measured the extent of heterogeneity of sequencing characteristics between droplet partitions. As the number of reads per genomic bin is approximately one, we hypothesized that the variation in coverage in a single bin across different cells would follow Poisson statistics. Hence, we plotted the coefficient of variation of each genomic bin’s coverage versus its mean. We observed a strong correlation between the coefficient of variation versus the mean as one would expect from a Poisson distribution **(Fig. 2*E***).

To determine CNVs for each cell, we modeled per-cell read counts per genomic bin as a Poisson distribution dependent on both the GC content and the copy number **(Methods)**. GC bias was modeled as a quadratic function with fixed intercept and correction on a cell-by-cell basis was performed. To estimate copy number for each bin, we empirically computed the effect of GC content **(Fig. *S*1*B*)** followed by scaling to generate haploid-scaled copy number calls. To identify candidate breakpoints, we calculated the discontinuity in copy number values among all mappable bins using the log-likelihood ratio statistic **(Fig. *S*1 *C*)**. Finally, read counts were centered on integer copy number states by numerical optimization to generate haploid-scaled calls. Full algorithmic and software details are available in **Supplementary Information**. The CNV calls were consistent with those produced by Ginkgo (11) **(Fig. *S*1*D*)**. CNVs were generally restricted to regions of the genome (88%) where reads could be confidently mapped **(Methods)**.

To assess the rate of false positive copy number calls, we calculated the copy number landscape of 1,046 cells of the PBMC control as described previously **(Fig. 2*F***). Copy number segments were divided into 20kb bins across the human genome for each cell. On average, less than 1% of 20kb bins in autosomal regions with high alignment score had a copy number other than 2. This result suggested a negligible false positive CNV calling rate. Specifically, aberrant calls occurred in regions of the human genome, which are difficult to align, such as centromeric and telomeric locations **(Fig. S1*E***).

### Determining Cell Cycle Status in Cancer Cells

We analyzed gastric cancer cell lines to determine replication- and aneuploidy-specific breakpoints as derived from WGS. We analyzed a total of 8,824 single cells from nine different gastric cancer cell lines **(Table S1, S2)**. We identified an average of 2,198 breakpoints per sample that were present in more than 1% of cells per cell line. The scDNA-Seq derived copy number and aneuploidy status was confirmed by SNP array analysis and karyotyping of these same cell lines **(Fig. S2)**. The average ploidy across cells was consistent with that reported by a separate karyotyping of these same cell lines **(Fig. S2*C*)**.

Difference in CNV signatures among cells is the result of subclonal populations with distinct copy number signatures or individual variation in cell cycle states. We focused our analysis of intratumoral heterogeneity on the subset of G0/G1 cells, to reduce the contribution of copy number changes attributable to a cell being in S phase **(Fig. 3*A*)**. For classifying cell cycle state we used three features: i) the cell’s ploidy, ii) its number of breakpoints and iii) the distance of breakpoints to replication origins (22). The proportion of G0/G1 cells ranged from 58% in SNU-16 to 82% in SNU-668 **(Fig. 3*B* and Table S1)**. For a subset of the cell lines, we used flow cytometry analysis to generate comparison data of DNA content **(Fig. S5*A*)**. The percentage of replicating cells per scDNA-Seq was positively correlated to the percentage of replicating cells per flow cytometry (r=0.86, P=0.063; **Fig. 3*C***). The percentage of G0/G1 cells per scDNA-Seq was also proportional to the doubling time of the cell line (r=0.76, P= 0.017; **Fig. 3*B***). Specifically, an extended duration in G0/G1 was an indicator of slower cell growth.

**Figure 3.**
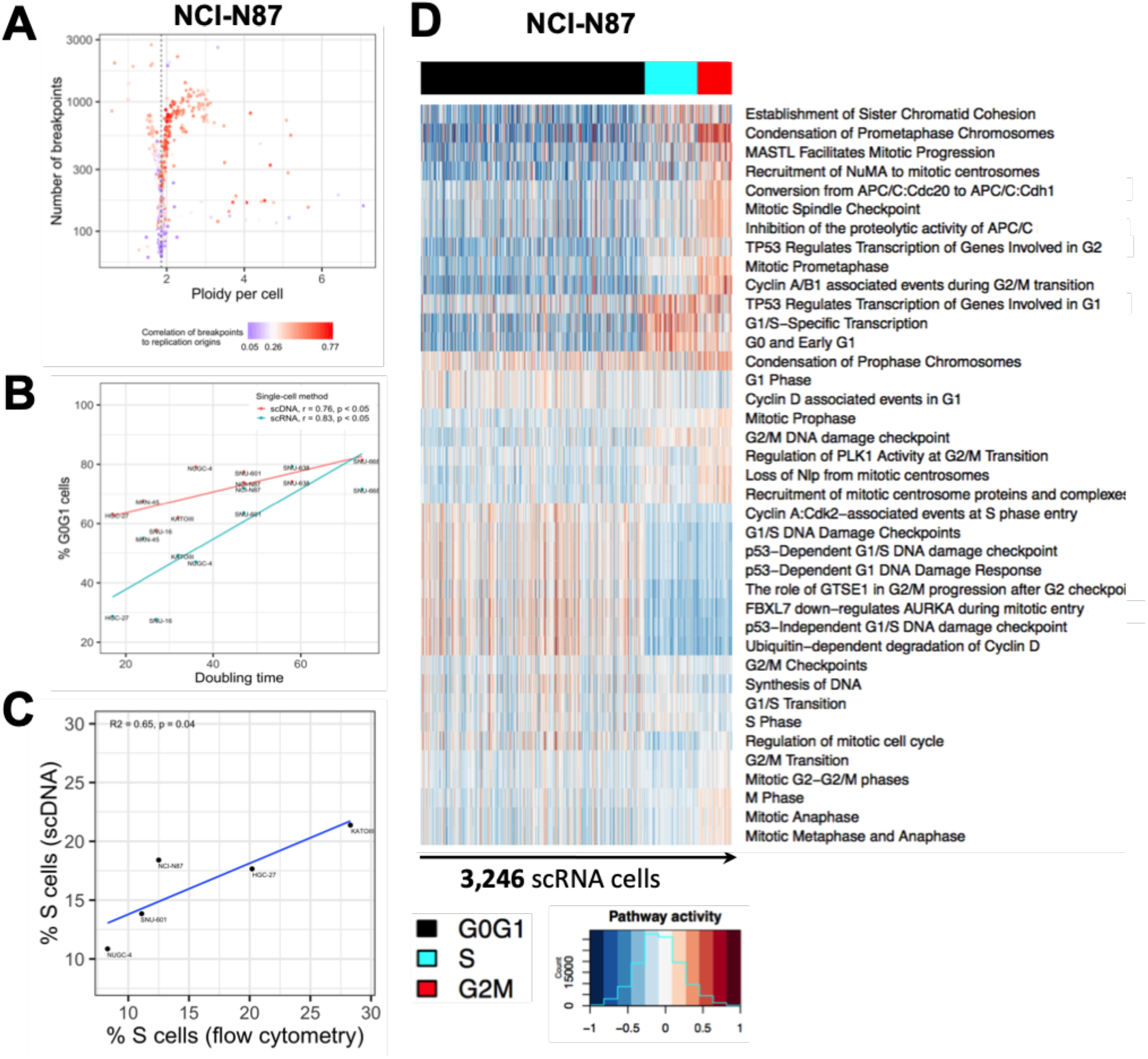
ScDNA- and scRNA-Seq delineate cell cycle state heterogeneity of gastric cancer cell lines. (**a-c**) scDNA-Seq derived cell cycle assignment. **(A)** 1,005 scDNA sequenced NCI-N87 cells are classified according to three features: their ploidy (x-axis), the number of breakpoints in their genome (y-axis) and their breakpoint’s proximity to human replication origins per chromosome (ORIs; color bar). Each cell’s uncommon breakpoints (i.e. breakpoints identified in <= 1% cells) are counted for each chromosome. For S-phase cells, these counts are correlated to the number of ORIs per chromosome. In contrast to S cells, G0/G1 cells have fewer breakpoints and their count is not correlated to chromosomal ORI counts. **(B)** % G0/G1 cells (y-axis) estimated from scDNA-Seq or scRNA-Seq is positively correlated with doubling time entries. **(C)** Validation of scDNA-Seq informed cell cycle phase assignment with flow cytometry. Cell lines shown were quantified by both techniques from the exact same suspension. **(D)** Cell cycle phase assignment of scRNA sequenced cells. 3,246 NCI-N87 cells (columns) are clustered according to the activity of 39 pathways related to various states along the cell cycle (rows). Clusters are then classified as containing either G0/G1 cells (black), cell in S-phase (cyan) or cells in G2M (red).

We used scRNA-Seq to validate our scDNA-Seq’s cell cycle assignment. We conducted scRNA-Seq of 28,209 single cells across the same nine gastric cancer cell lines **(Table S3)**. Differences in passage number between scDNA- and scRNA-Seq experiments were kept to a minimum and the extent of confluence was typically at 80-90% **(Table S2)**. Activity profiles of multiple cell cycle pathways have been shown to provide robust cell cycle status classification across different cell types (23). For each individual cell, we quantified the activity of 39 cell cycle pathways from the REACTOME database (24) and used these results to determine cell cycle state **(Table S4)**. Pathways were classified into three groups depending on their main activation timing during G0/G1, S, and G2M. We performed hierarchical clustering of cells and classified clusters based on their cells’ pathway activity **(Fig. 3*D*)**. The percentages of G0/G1 cells, assigned with scDNA-Seq versus scRNA-Seq were highly correlated (r=0.73, P=0.026, **Fig. 3*B***).

### Subclonal Signatures of Genomic Instability and Ongoing Selection with scDNA-Seq

We used scDNA-Seq to characterize the underlying subclonal structure of the cell lines. Approximately 95% of the breakpoints identified by scDNA-Seq were found in less than 1% of the G0/G1 population – we ascribed these events to variance related to DNA replication and not representing true cancer CNVs **(Table S1)**. We used the remaining CNV segments and patterns of genomic instability to identify subclones within the G0/G1 population. Using the CNV features, we calculated the pairwise distances between cells in Hamming space, thus assigning a higher weight to larger genomic segments. We applied a neighbor joining algorithm, BIONJ (25), to build a phylogenetic tree of G0/G1 cells **(Methods)**. We defined a clone as the largest subtree within which the maximum distance between its cell members was less than 20% of the affected genome **(Fig. 4*A*)**. The relative fraction of cells assigned to a subclone is referred to as subclone size. To assign S-phase cells to the subclones detected among the G0/G1 population, we determined the cellular similarity with a Pearson correlation. For example, this approach identified four clones within the G0/G1 population of NCI-N87 **(Fig. 4*A,B***). The percentage S cells assigned to each of these four subclones were proportional to their respective G0/G1 representation **(Fig. 4*A,B*)**.

**Figure 4:**
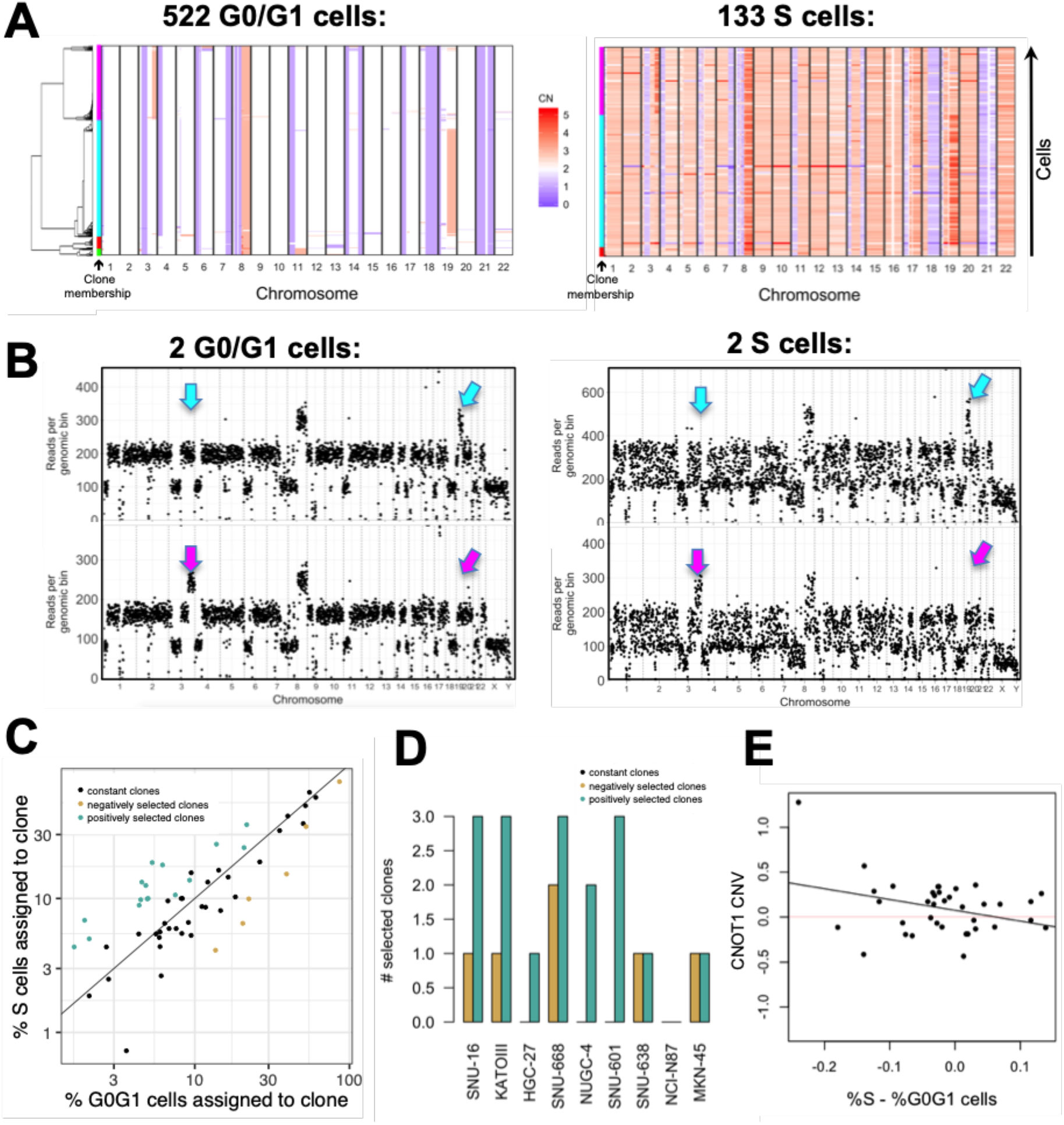
Intra-tumor heterogeneity and evolution in gastric cancer cell lines. **(A)** Copy number landscape of G0/G1 cells (left) is shown alongside S cells (right) for each clone detected in NCI-N87 (left color bars). **(B)** Copy number segmentation profile shown for an G0/G1- and an S representative of the two largest clones in (a) (cyan and purple). Arrows indicate genomic regions where the two clones diverge. **(C)** % Replicating cells per clone increases with % G0/G1 cells per clone in NCI-N87 as well as in the other eight cell lines, indicating clonal stasis (Pearson r = 0.88; p < 2e-16). Selection of clones (color-coded) calculated as probability of sampling the % replicating cells observed for a given clone, conditional on the G0/G1 representation of that same clone using the hypergeometric distribution (**Methods**). Clones are assigned to three groups: positive selection (n=17), no selection (n=34) and negative selection (n=6). **(D)** Number of selected clones per cell line. **(E)** Copy number of mRNA deadenylase *CNOT1* relative to baseline ploidy (y-axis) is higher among clones with lower selection coefficients (x-axis). Selection coefficient calculated by subtracting the % G0G1 cells from the % S cells assigned to a clone.

Based on our analysis across the gastric cancer cell lines, anywhere from two to 12 subclones were present per a cell line **(Table S5)**. Approximately half of the variation in subclones per cell line was attributed to the cell lines’ ploidy and/or the duration since the cell line was first established in culture (adjusted R^2^=0.53; p=0.044; **Table S6**). Higher ploidy predicted more clones (coefficient = 3.09; p = 0.054), while longer time in culture was predictive of fewer clones (coefficient = −0.29; p = 0.025). The latter observation is consistent with a recent finding showing that *in vitro* CNV acquisition rate decreases over time, while signatures of proliferation increase, in line with clonal selection of fitter clones (9).

Shifts in a cancer cell line’s subclonal composition have been shown to frequently result from *in vitro* selection, rather than stochastic processes (9, 10). To quantify in vitro selection among the cancer cell lines, we compared the percentage replicating cells per subclone to the percentage G0/G1 cells in that subpopulation. The two cell cycle states had similar proportions for a given subclone, indicating predominance of clonal stasis (Pearson r = 0.88; p < 2e-16; **Fig. 4*C***). We used a hypergeometric distribution to model the subclone’s number of replicating cells and test if it was within a range consistent with its G0/G1 representation ***(Supplemental Methods***). Seventeen subclones (30%) had a higher percentage of replicating cells than expected from their G0/G1 population size (FDR adjusted P<=0.05; **Fig. 4*C,D***). We interpreted this result to be a possible indication of positive selection. Conversely, six subclones (11%) had fewer replicating cells than expected from their G0/G1 representation, suggesting they were under negative selection (FDR adjusted P<=0.05; **Fig. 4*C,D***).

We found that negatively selected subclones were enriched for a CNV gain of *CNOT1* - a gene involved in mRNA degradation **(Fig. 4*E***), and for deletions of *IREB2, SIN3A* and *MAP2K1* (|Pearson r| >= 0.32; p<=0.05). The overrepresentation of positively selected clones compared to negatively selected ones was consistent with a recent study showing that *in vitro* evolution is primarily driven by positive selection (10). Quantifying the selection of individual subclones may prove useful in predicting the genome states of future cell line populations.

### Consilience of scDNA- and scRNA-Seq on G0/G1 subclonal architectures

We demonstrated scDNA-Seq provides high resolution analysis of CNVs than single cell gene expression studies. In addition, the use of joint sets offered a way of determining comparison of clonal overlap. First, we determined whether scRNA-Seq identified the same set of subclones derived from scDNA-Seq. For this comparison, we inferred CNVs from scRNA-Seq. Gene expression has been shown to be proportional to the gene’s copy number state for the majority of genes (26), suggesting that scRNA-Seq derived expression features can inform CNV status. However, other mechanisms of gene regulation alter expression, confounding the influence of segmental copy number. This is most evident when analyzing short genomic segments, below 10 Mb. One algorithm for calling CNVs from scRNA-Seq data demonstrated good performance, particularly for large segments, above 10 Mb, and for large subclones (27). However, this method’s precision fell below 50% for smaller subclones, making up 20% or less of the total cells (27). To address this issue, we developed and applied an algorithm called LIAYSON, which uses scRNA-Seq to deconvolute bulk CNV profiles into single cell specific copy numbers ***(Supplemental Methods***). This approach relies on gene expression to estimate the variance in copy number, but not the mean copy number across cells (**Fig. S3*A,B***), and is less influenced by regulators of expression levels other than CNVs. It requires on average at least 20 genes with expression results from a genomic interval of at least 10 Mb. Between 25-80% of segments per cell line passed these metrics for this analysis.

With the CNV results from scRNA-Seq, we identified a range of three to 11 subclonal populations across the nine gastric cancer cell lines **(Table S5)**. The number of scRNA-Seq and scDNA-Seq derived clones were highly correlated (r=0.93. P=3E-4; **Fig. 5*A***). As another validation of concordance between the two methods, we performed hierarchical clustering of subclone-specific CNV profiles **(Fig. 5*B-D*)**. We defined true positives as clusters containing subclones identified by both methods and false positives and negatives for clusters containing only one type of subclones but not the other ***(Supplemental Methods***). The concordance between scDNA-Seq and scRNA-Seq had an average F1 score of only 0.47 for clones below 4% abundance, but increased to >=0.7 for clones above 4% size **(Fig. S3*C*)**. Based on this result, we excluded any subclones smaller than 4% size and not confirmed by both single cell methods. Posteriori saturation curves of scDNA-Seq library sizes were calculated for each cell line as previously described (2) and indicated that we had statistical power to detect these subclones **(Table S7)**.

**Figure 5.**
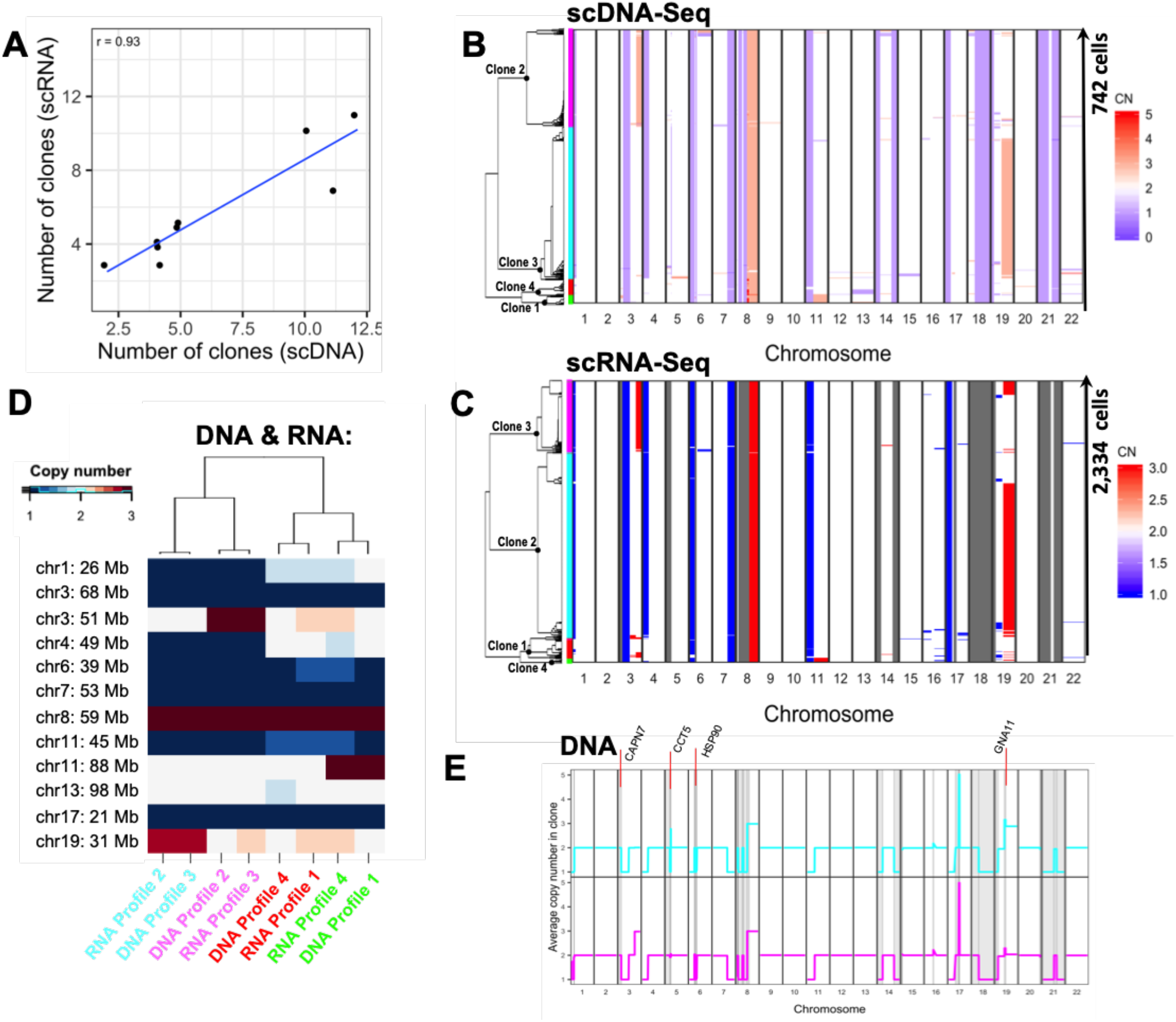
Consilience of scDNA- and scRNA-Seq on G0/G1 clonal architectures. **(A)** Correlation between number of clones inferred by scRNA- and scDNA-Seq. **(B)** ScDNA-Seq derived copy number landscape (columns) of 742 G0/G1 cells (rows) distinguishes four clones. Clone membership is color coded on the left. **(C)** scRNA-Seq derived copy number landscape of 2,334 G0/G1 cells independently distinguishes four clones (left color bar). **(D)** Moreover, each CNV profile found by scRNA-Seq has an equivalent CNV profile in the scDNA-Seq data, applying to a similar % of cells. **(E)** Clone specific differences in CNVs are shown for the two largest NCI-N87 clones (purple and cyan) along with affected cancer genes in those regions. Highlighted as gray bands are genomic regions too small to be assigned clone-specific CNVs by scRNA-Seq and thus detectable only with scDNA-Seq.

Citing an example, our scDNA-Seq and scRNA-Seq results identified four subclones in NCI-N87 with similar proportional sizes **(Fig. 5*B-D*)**. On closer examination we observed that the copy number states of several smaller segments (<10 Mb), were not assigned for any clone by scRNA-Seq, but were identified by scDNA-Seq. For these genomic regions, the number of genes with adequate expression levels was too low to allow assignment by scRNA-Seq **(Fig. 5*E*)**. Therefore, our conclusion was that scDNA-Seq provided higher degree of subclonal characterization.

The other gastric cancer cell lines had similar results. Among the nine cell lines, the subclonal size, as determined by scRNA-Seq, correlated with the scDNA-Seq results (Pearson r=0.93, p < 2e-16; **Fig. S3*D***). An average of 88% cells per cell line were assigned to subclones confirmed by both scDNA-Seq and scRNA-Seq **(Table S5)**. The discordance was attributable to subclones with a 10% or lower cellular fraction. Concordance between the two methods was dependent on the sequence depth, the subclone size **(Fig. S3*C-F*)** and the subclonal number per a given cell line **(Table S5)**. For SNU-16 and SNU-668, differences in passage number between scDNA- and scRNA-Seq experiments were correlated with a greater divergence between clonal compositions measured by the two methods (r=0.71, P=0.032; **Fig. S3G)**. Thus, the CNV comparisons demonstrated subclonal overlaying with scDNA-Seq providing higher resolution of subclonal genomic characteristics.

### Analyzing a Primary Gastric Cancer with Joint Single Cell Sequencing

As a test of joint single cell genomics on a clinical tissue sample, we analyzed a Stage II gastric adenocarcinoma **(P5931)**. We determined CNVs, gene expression, subclonal assignment and pathway activity for a given single cell. Histopathology of this gastric cancer revealed moderate to poorly differentiated features with a 60-70% tumor fraction. Immunohistochemistry demonstrated a loss of *MLH1* and *PMS2* expression. The loss of these proteins indicated that this tumor had microsatellite instability **(MSI)** where cancer cells have a hypermutable state because of loss of DNA mismatch repair. The tumor tissue was disaggregated into a single cell suspension and analyzed with both single cell genomic methods **(Methods)**.

From this patient-derived tumor sample, we sequenced 796 cells using scDNA-Seq and 2,098 cells using scRNA-Seq. The G0/G1 representation was 79.5% among scDNA- and 61% among scRNA- sequenced cells **(Fig. S4*A,B*)**. After eliminating cell-cycle related breakpoints **(Fig. S4*A*)**, a total of 28 CNVs were identified among the G0/G1 population, including gain of chromosome 8, 3 and 19q – three of the four allelic imbalances are among the most common events in MSI-positive gastric cancers (28). Both scDNA-Seq and scRNA-Seq identified four clones **(Fig. 6*A*)** with three being concordant and making up over 90% of the cells in each assay. A diploid subpopulation comprised 59% of the scDNA- and 50% of the scRNA-sequenced G0/G1 population **(Fig. 6*A*)**.

**Figure 6.**
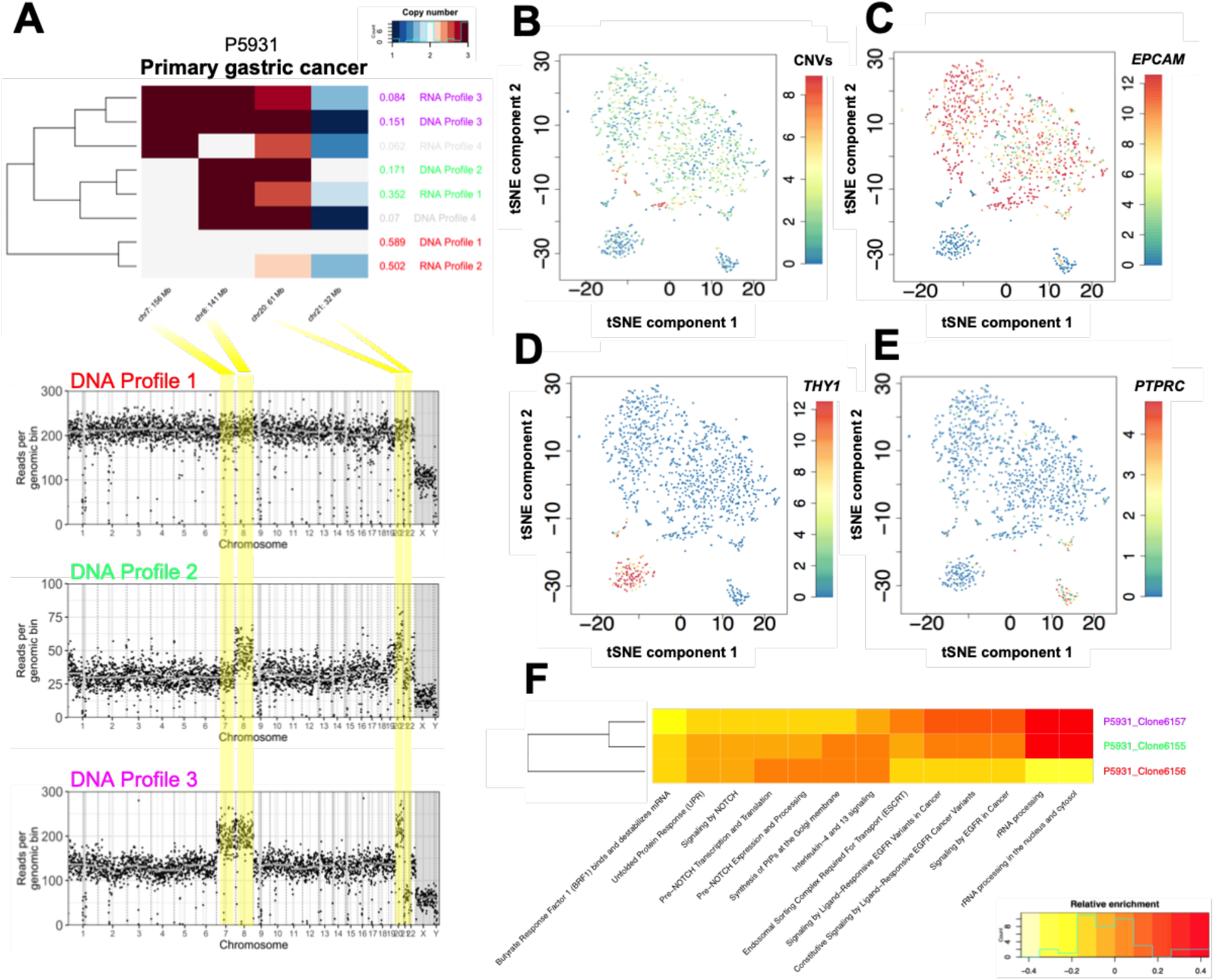
Integrated scDNA- and scRNA-Seq analysis of patient 5931’s G0/G1 population. **(A)** Both scRNA- and scDNA-Seq independently identify four clones in P5931, each with a distinct CNV profile. Three out of four CNV profiles found by scRNA-Seq have an equivalent CNV profile in the scDNA-Seq data, applying to a similar % of cells. Clone membership is color-coded on the right (gray – identified by either scDNA-Seq or scRNA-Seq alone; rainbow – confirmed by both techniques). Copy number segmentation profiles are shown for three cell representatives – one for each of the three confirmed clones. Yellow bands indicate genomic regions where the clones diverge. (**b-e**) TSNE map of 1,090 G0/G1 cells calculated based on the expression of variable genes (**Methods**). Non-epithelial cells are marked by absence of CNVs **(B)** and of *EPCAM* expression **(C)**, and include myofibroblasts – identified by THY1 **(D)**, and immune cells – identified by higher levels of PTPRC expression **(E)**. **(F)** Differential activation of pathways among epithelial representatives of the three confirmed clones (Anova: P<0.001).

The scRNA-Seq results provided transcriptional features for each subclonal population. Consistent with immunohistochemistry, *MLH1* expression was mostly absent, but *PMS2* expression was low to moderate (detected in 1.6% and 16.5% of G0/G1 cells respectively). To distinguish the non-epithelial population from epithelial, we used differences in the epithelial marker *EPCAM.* In contrast to *EPCAM*- cells (27% of the G0/G1 population), which were diploid, the majority of *EPCAM+* cells harbored CNVs **(Fig. 5*B,C*)**. Thus, most of the epithelial cells were cancer, while most diploid cells were non-epithelial. After excluding the myofibroblasts/fibroblasts (13%), endothelial cells (1.6%) and immune cells (12.3%) **(Fig. 5*D,E*)**, the normal, diploid subpopulation comprised 47% of the epithelial population. Differences in the activities of canonical pathways among the various epithelial cellular subpopulations (i.e. tumor, normal) were calculated with Gene Set Variation Analysis **(GSVA)**. The top pathway functions enriched in the tumor epithelial cells compared to normal epithelial cells included upregulation of genes involved in epidermal growth factor receptor **(EGFR)** signaling **(Fig. 6*F* and Table S8)**.

On close examination, differential pathway activities were apparent when comparing two subclones **(Fig. 6*A*)**. Compared to the larger of the two subclones **(Fig. 6*A*)**, the smaller one had a higher Ras activation pathway changes and increase expression of genes involved in biotin transport (**T-test: P<0.005**), that suggested higher metabolic activity **(Fig. 6*F*)**. In contrast, pathways indicating Notch signaling and laminin interactions had lower transcriptional levels in the smaller tumor subclone (**T-test: P<0.005**). Measuring the subclone specific activity of pathways can thus inform hypotheses on the importance of different phenotypes indicative of biological divergence during early versus late stages of tumor evolution.

## DISCUSSION

For this study we demonstrated a new scDNA-Seq technology that enabled the interrogation of intratumoral heterogeneity from thousands of cells per sample. We demonstrated how a joint analysis, adding RNA-Seq at the resolution of single cells, provided additional supporting information about the characteristics of cancer evolution in the context of cellular heterogeneity. In contrast to a prior study, that joined scDNA-Seq with scRNA-Seq to identify subclones (3), we chose to compare subclones identified independently by each single cell technology, improving subclone validation. For this aspect of the study, we developed a new method that leverages association-rule mining to infer large-scale CNVs from scRNA-Seq. Co-clustering clones identified by either single cell method intrinsically controlled for false positives: whether two clones co-cluster not only depends on their own genetic content, but also on the content of other clones identified in the sample. We demonstrated that this type of joint analysis can be conducted on primary tumor biopsies.

Integrating the transcriptome and genome features allowed us to characterize the genetic basis of clonal expansions and identify seminal features of underlying pathway dysregulation across diverse clonal populations. Subsequently, we demonstrated that this analysis’ utility on primary tumor biopsies. In the future, clone specific marker candidates such as surface proteins, could inform flow cytometry sorting of clones to study their differential drug sensitivities.

Our study showed that gastric cancer cell lines have substantial genetic subclonal diversity. This result is consistent with other studies showing that cancer evolution continues *in vitro* (9, 29). Cell line heterogeneity has implications for in vitro drug studies where the clonal composition may prove to be an important factor. Moreover, one can use cell lines for in vitro studies of clonal competition, therapeutic adaptation and transcriptional reprogramming (3). Cellular diversity in cancer cell lines can be the result of stochastic drift or of ongoing selection during tissue culture passaging. Several approaches have been developed to quantify selection using either time-resolved sequence data from longitudinal studies (30, 31) or by observing differences in the statistical structure and shape of genealogies reconstructed from a fitness diverse asexual population (32, 33). Our integrated sequencing approach enabled prediction of selection strength by simply comparing S to G0/G1 representations of a clone. Our study demonstrated results consistent with behavior demonstrating tumor evolution where a significant proportion of subclones undergoing both positive and negative selection. This result is consistent with prior observations that cancer cell line diversification is a consequence of *in-vitro* selection (10). In the future, the coexistence of multiple clones within the same cell line can be leveraged to learn generally applicable strategies that differentiate between the sensitivities of co-existing clones and that characterize clonal competition, cooperation, and the effect of diversity on adaptation. For future studies, we will use joint single cell analysis for both translational studies of clinical tumor samples as well as *in vitro* studies to relate a cell’s level of genomic instability to its fitness relative to cancer therapy.

## MATERIALS AND METHODS

### Cell lines and gastric cancer patient sample

Gastric cancer cell lines were purchased from ATCC (KATOIII, NCI-N87, SNU-16), KCLB (SNU-668, SNU-601, SNU-638), JCRB (MKN-45, NUGC-4) and ECACC (HGC-27). Identity of cells were determined through independent karyotyping. Cells were checked for mycoplasma contamination. Cells were cultured in their recommended media conditions at 37°C. Afterwards, the cells were processed into suspensions with standard procedures. Briefly, this process involved trypsinizing the cells, followed by inactivation by FBS. We performed washes by centrifugation at 400g in 1X PBS with 0.04% BSA. To remove cellular debris and cellular aggregates, we filtered cells through a Flowmi cell strainer (Wayne, NJ) before proceeding to single-cell DNA and RNA sequencing.

For analysis of clinical tumor samples, our study was approved by the Institutional Review Board at Stanford University. Informed consent was obtained from the patient. Tissue biopsies were obtained from surgical resection of a primary gastric adenocarcinoma and matched adjacent normal tissue. Immediately after resection, the tumor sample was stored in RPMI medium on ice for less than 1 hour. The sample was then macrodissected and dissociated into a cellular suspension by the gentleMACS Octo Dissociator using the human tumor dissociation kit as per manufacturer’s recommendations and the 37C_h_TDK_3 program (Miltenyi Biotec, Bergisch Gladbach, Germany). The suspension was used immediately for scRNA-seq. Single cell DNA-Seq was performed after thawing cryopreserved sample stored in liquid nitrogen in 90% FBS-10% DMSO freezing medium.

### Library preparation protocol for scDNA-Seq

Single-cell DNA libraries were generated using a high-throughput, droplet-based reagent delivery system using a two-stage microfluidic procedure. First, cells were encapsulated in a hydrogel matrix and treated to lyse and unpackage DNA. Second, a gel bead (GB) was functionalized with copies of a unique droplet-identifying barcode (sampled from a pool of ~737,000) and co-encapsulated with the hydrogel cell bead in a second microfluidic stage to separately index the genomic DNA (gDNA) of each individual cell. Unless otherwise stated, all reagents were part of a beta version of the Gel Bead and Library Kit for single cell CNV analysis (10X Genomics Inc., Pleasanton, CA).

In the first microfluidic chip, cell beads were generated by partitioning approximately 10,000 cells of each sample in a hydrogel matrix. A cell suspension is combined with an activation reagent, hydrogel precursors, paramagnetic particles, and loaded into one inlet well. In the other two inlet wells, CB polymer reagent and partitioning oil were added. To ensure a low multiplet rate, cells were delivered at a dilution such that the majority of CBs contain either a single cell or no cell. Once generated, the emulsion was immediately transferred into a PCR strip tube and incubated with orbital shaking at 1000 rpm overnight. The incubation yields polymerized magnetic CBs for subsequent steps.

Encapsulated cells were processed by the addition of lysis and protein digestion reagents to yield accessible DNA for whole-genome amplification. The presence of magnetic particles in the cell bead matrix enabled CB retention and streamlined washing and buffer exchange steps. After lysis, CBs were washed by magnetic capture, concentrated by reduction of liquid volume, and buffer exchanged with the addition of 1X PBS buffer. CBs were then denatured by NaOH, neutralized with Tris, and diluted in storage buffer. Finally, aggregates of cell beads were removed by filtration through a Flowmi strainer before a volume normalization procedure to set the CB concentration.

Cell bead-gel bead were generated by loading CBs, barcoded gel beads, enzymatic reaction mix, and partitioning oil in a second microfluidic chip. A majority of the **CBGBs** (~80%) contained a single CB and a single gel bead, which once encapsulated then dissolved to release their contents. To amplify and barcode gDNA, the emulsion was then incubated at 30°C for 3 hours, 16°C for 5 hours, and finally heat inactivated at 65°C for 10 minutes before a 4°C hold step. This two-step isothermal incubation yielded genomic DNA fragments tagged with an Illumina read 1 adapter followed by a partitionidentifying 16bp barcode sequence.

The emulsion was broken and purified as previously described^1^. Conventional end-repair and a-tailing of the amplified library was performed, after which a single-end sequencing adapter containing the Illumina read 2 priming site was ligated. PCR was performed using the Illumina P5 sequence and a sample barcode with the following conditions: 98°C for 45 seconds, followed by 12-14 cycles (dependent on cell loading) of 98°C for 20 seconds, 54°C for 30 seconds, and 72°C for 30 seconds. A final incubation step at 72°C was performed for 1 minute before holding at 4°C. Libraries were purified with SPRIselect beads (Beckman Coulter, Brea, CA) and size-selected to ~550bp. Finally, sequencing libraries were quantified by qPCR before sequencing on the Illumina platform using NovaSeq S2 chemistry with 2×100 paired-end reads.

### ScDNA-Seq data processing and CNV calling

Sequencing data was processed with the Cellranger-dna pipeline, which automates sample demultiplexing, read alignment, CNV calling, and report generation. In this study, we used a beta version for all analyses (6002.16.0). Paired-end FASTQ files and a reference genome (GRCh38) are used as input. Cellranger-dna output includes copy number calls for each cell.

The computational pipeline includes **preprocessing** and **single cell copy number calling.** The outputs of this pipeline are CNV calls *and* read counts in 20kb bins across the genome as genomic bin- by-cell matrices. A summary is provided in this section – full details are included in the **Supplementary Material**. In the preprocessing stage, the first 16 base pairs of read 1 are compared to a whitelist of all possible droplet barcodes (totaling ~737,000). All observed droplet barcodes were tested for the presence of a cell by using mapped read abundances to the human genome. Reads were aligned to GRCh38 using bwa-mem. Each read in the bam file was annotated with a cellular barcode tag ‘CB’. Confidently mapped reads were counted across the genome in 20kb non-overlapping windows. GC bias correction, modelled as a polynomial of degree 2 with fixed intercept, was applied. Copy number calls are determined by modeling binned read abundances to a Poisson distribution with the copy number, GC bias, and a scaling factor as parameters. Candidate breakpoints were estimated by applying a log-likelihood ratio statistic against fluctuations in read coverage over neighboring genomic bins. These breakpoints were refined and reported as a set of non-overlapping segments across the genome. The copy numbers were scaled to integer-level ploidies. Copy number calls for non-mappable regions were imputed with neighboring copy number calls in confidently mapped regions, provided that the copy number on both sides of a non-mappable region were the same and the region was < 500 kb.

### LIAYSON: Calling CNVs from scRNA-Seq

**Li**nking single-cell genomes **a**mong contemporar**y s**ubcl**one** transcriptomes **(LIAYSON)** is an approach to profile the CNV landscape of each scRNA-sequenced single cell of a given sample. The algorithm relies on two assumptions: i) a cell’s average copy number state for a given genomic segment influences the mean expression of genes within that segment across the same set of cells; and ii) the copy number variance of a given genomic segment across cells reflects the cells’ expression heterogeneity for genes within that same segment ***(Fig. S*3A,B)**. LIAYSON is available at the following URL https://github.com/noemiandor/liayson.git.

LIAYSON’s algorithm involves the following: Let x’ ∈ CN be the measured copy number of a given cellsegment pair, and x its corresponding true copy number state. The probability of assigning copy number x to a cell i at locus j depends on: a) cell i’s read count at locus j and b) cell i’s read count at other loci, i.e. how similar the cell is to other cells that have copy number x at locus j. For (a), we fit a Gaussian kernel on the read counts at locus j across cells to identify the major and the minor copy number states of j as the highest and second highest peak of the fit respectively (**Supplementary Methods**). For (b), we use Apriori (34) – an algorithm for association rule mining – to find groups of loci that tend to have correlated copy number states across cells (**Supplementary Methods**). LIAYSON is implemented in R and is available on CRAN.

### Identification of coexisting clones from scDNA-Seq or scRNA-Seq

Let CNF be the matrix of copy number states per non-private segment per G0/G1 cell, derived either from scRNA- or from scDNA-Seq, with entries (i, j) pointing to the copy number state of cell i for segment j. Pairwise distances between cells were calculated in Hamming space (35) of their segmental copy number profiles (rows in CNF), weighted by segment length. We used the BIONJ algorithm (25) to reconstruct a phylogenetic tree of G0/G1 cells from the distance matrix. A subtree was defined as a clone if the maximum distance between its cell members was less than 20% of the genome. Finally, we used the Pearson Correlation Coefficient calculated across segments to assign S and G2M cells to the clones detected among the G0/G1 population. The copy number profile of each detected subclone was calculated as the average profiles of assigned subclone members.

### Integration of scRNA-Seq- and scDNA-Seq derived clones

Let R and D be the scRNA- and scDNA-Seq derived clone-by-segment matrices of copy number states. Furthermore, let S:= S_R_ ⋂ S_D_, where S_R_ and S_D_ are the segments defining the columns of R and D respectively. We defined X:= R_S_ U D_S_ which was the union of scRNA-Seq and scDNA-Seq derived clones at overlapping genomic locations. We used the same hierarchical clustering procedure as above, only this time clones rather than cells were arranged into the resulting dendrogram T. We iterated through all binary subtrees t ∈ T and assigned clones within t as:

i. True positives (TPs) – t contains both, an scRNA- and an scDNA-clone
ii. False positives (FPs) – t contains two scRNA-clones
iii. False negatives (FNs) – t contains two scDNA-clones.

To validate scDNA-Seq derived clone detection, we used the same procedure, except the roles of FPs and FNs were flipped. Clones comprising less than 4% cells, which were not confirmed by both techniques, were excluded from further analysis.

## Author’s contributions

N.A., B.T.L., A.J., C.C., V.K., K.B., T.D.W., A.D.P., M.S., R.W.D., R.B. and H.P.J. contributed to the experimental design. C.C., V.K., K.B., T.D.W., A.D.P., M.S., Z.D., J.S., Z.B., B.K.L., A.J.M., M.R., S.S.S., M.S., J.S., K.S., A.W., W.Y., Y.Y., B.T.L., M.S-L., R.B. contributed to the technology development. B.T.L., A.S., J.C., C.C., M.A.K., T.D.W., A.D.P., M.S., L.D., S.J., B.K.L., A.J.M., M.R., S.S.S., M.S., J.S., K.S., A.W., W.Y. and Y.Y. conducted the experiments. V.K., D.S., L.H., N.A. and B.T.L. contributed to the software development. N.A., B.T.L., C.C., V.K., K.B., S.M.G., T.D.W., A.D.P., M.S., L.H., S.M., M.S-L., R.B. and H.P.J. contributed to the data analysis. C.K.S., G.A.P. and H.P.J. contributed to the clinical sample analysis. N.A., B.T.L., V.K. and H.P.J. did the writing. R.B. oversaw the technology development and CNV calling software. H.P.J. oversaw all aspects of the genomics studies.

## Availability of data and material

The datasets generated for this study are available in the National Institute of Health’s dbGAP repository; accession number phs001711. LIAYSON is available at the following URL https://github.com/noemiandor/liayson.git.

## Acknowledgements

This work was supported by the following grants from the NIH: NHGRI P01HG000205 to BTL, MAK, SMG, RWD and HPJ, NCI K99 CA215256 to NA, NHGRI R01HG006137 to SMG. and HPJ. and NCI U01CA217875 to AS and HPJ. The American Cancer Society provided additional support to JS and HPJ [Research Scholar Grant, RSG-13-297-01-TBG]. In addition, HPJ received support from the Clayville Foundation, the Gastric Cancer Foundation and the Seiler Family Foundation. We thank Dino Valdecanas for assistance with chip drafting and Minji Kim for assistance with chip development. Also, we thank Michelle Luo and Steven Short for reviewing the manuscript.

