## Supplemental Tables, Figures and Methods for "Joint single cell DNA-Seq and RNA-Seq of cancer reveals subclonal signatures of genomic instability and gene expression"

**INSTITUTIONS**

[1]Division of Oncology, Department of Medicine, Stanford University School of Medicine, Stanford, CA, United States

[2]Stanford Genome Technology Center, Stanford University, Palo Alto, CA, United States

[3]10X Genomics, Pleasanton CA, United States

[4]Department of Surgery, Stanford University School of Medicine, Stanford, CA, United States

[5]Department of Pathology, Stanford University School of Medicine, Stanford, CA, United States

**TO WHOM CORREPSONDENCE MAY BE ADDRESSED**

Hanlee P. Ji

Rajiv Bharadwaj

**TABLE OF CONTENTS**

| | |
|---|---|
| Supplementary Methods – Overview of CNV pipeline ('cellranger-dna') | Pages 1 - 6 |
| Supplementary Methods – LIAYSON: Calling CNVs from scRNA-Seq | Pages 7 - 8 |
| Supplementary Figures | Pages 9 – 14 |
| Supplementary Tables | Pages 15 - 22 |

**SUPPLEMENTARY METHODS – Overview of CNV pipeline (‘cellranger-dna’)**

***Preprocessing - alignment:*** The first 16 bases of read 1 in each read pair is a droplet-labeling barcode. The 16-base barcode can take one of 737,000 different sequences that comprise a whitelist. Due to sequencing error the observed barcode may not be a perfect match to whitelist barcode. When a raw barcode is one mismatch away from a whitelisted barcode we performed error correction as previously described[1,2]. We aligned all the trimmed read pairs (trimmed read 1, read 2) to the genome using bwa-mem with flag -M. The alignment procedure produced a bam file where each read is labeled with a CB tag that contains the error corrected barcode sequence followed by a number, e.g., CB:Z:CTACCCAAGTCGACTT-1.

***Preprocessing - genome mappability:*** We restricted copy number calls to regions where sequence reads confidently mapped to the reference genome of interest. We used an empirical method to determine the reference mappability that is specific to a sample’s sequence data. The mappability of each bin was calculated by simulating reads from the reference genome and measuring the proportion of reads mapping to their expected location.

Let $L1$, $L2$ denote the number of bases sequenced on read 1 and read 2 respectively. We simulated perfect reads from the reference genome at 1X coverage where read 1 and read 2 have lengths $L1-16$ and $L2$ respectively and an insert size distribution that mirrors the input library. We map the simulated reads back to the reference genome using the same procedure described above to map the sequencing reads. We divided each chromosome of the genome into 20 kb bins. For each genome bin $\alpha$ we computed the fraction $m_\alpha$ of simulated reads from that bin that map back to it with mapping quality $\geq 30$. This defines the mappability across the genome. Bins composed entirely of *N* bases would have mappability = 0, bins composed of sequences that are unique in the genome will have mappability = 1.0; and bins with highly repetitive content or bins that lie within duplications in the reference genome will have low mappability. In addition, we computed the GC composition in each 20kb bin, which is denoted by $g_\alpha$. For the human reference genome GRCh38 and a library sequenced to 2x100 about 88 % of genome bins have a mappability > 90 %.

***Preprocessing – cell detection:*** Droplet barcodes labeled the partitions where the barcoding reaction takes place, but not all partitions contains cells. The reads per barcode distribution for an experiment is shown in **Fig. 1A**. Cells, numbering *N* in total for an experiment, are defined by the following procedure:

1. Determine the barcode associated with the highest number of reads = *M*.
2. Construct the set of barcodes $B_0$ where each barcode is associated with at least $M/10$ reads.
3. Compute $M_{99}$ = the 99th percentile of the set $B_0$.
4. Barcodes with at least $M_{99}/10$ reads are defined as cells.

***Pre-processing – coverage profile matrix:*** We divided each chromosome of the reference genome into 20 kb bins. For simplicity of representation we considered the concatenated genome: a single vector $G_\alpha$ for $\alpha = 0,\ldots,L_G$, where $L_G$ is the number of 20 kb bins in the genome. We organized the aligned reads by barcode and computed the coverage per cell per 20kb bin matrix denoted by $X$. Each element in $X$, $X_{i\alpha}$, represents the number of read-pairs in cell $i$ in bin $\alpha$. We only consider read pairs that are not duplicates and have mapping quality ≥ 30. Duplicate read pairs are marked using the start and stop position of the pair alignment, and read pairs with distinct cell barcodes are never marked as duplicates of each other. Reads contribute individually to $X$: aligning reads each contribute 0.5 to $X_{i\alpha}$, except in the cases of an unmapped mate (contributing 1.0), and where the mate maps beyond 20kb or to a different chromosome (contributing 1.0).

***Copy number determination:*** The process of calling copy number of a single cell (or a group of similar cells) starts from the read coverage. Our notation is as follows (some already described in earlier sections):

- $x_\alpha$ denotes the read coverage (a single row of the coverage matrix X defined above).
- $g_\alpha$ denotes the GC content per bin.
- $m_\alpha$ denotes the mappability per bin.
- $\alpha = 0,\ldots,L_G - 1$ is the bin coordinate along the genome.
- $\tilde{\alpha} = 0,\ldots,L_{GM} - 1$ is the bin coordinate along the genome ignoring all bins with mappability ≤ 0.90, where $L_{GM}$ is the total number of bins with mappability > 0.90.
- $p_\alpha \in \mathbb{Z}$ denotes the copy number per bin.
- $\sigma$ is the number of reads that a copy number 1 (haploid) 20 kb DNA fragment would produce on average. This is a measure of the efficiency of conversion of genomic DNA into sequenced read pairs and we simply call this the scale.

We modeled $x_\alpha \sim \text{Poisson}(\sigma p_\alpha f(g_\alpha))$, where $f$ is an arbitrary positive function over the domain [0,1]. The problem at hand is to infer $p_\alpha$ starting from the observed read counts $x_\alpha$ and the GC content $g_\alpha$. Our strategy involves first computing the effect of GC content, $f(g_\alpha)$ and then estimating the scale $\sigma$.

Note: we ignore regions of the genome where the mappability per 20kb bin is ≤ 0.90. For the human reference genome GRCh37 and a typical 10x single cell DNA library sequenced to 2x100 bp reads this translates to about 87% of the reference genome. We perform copy number calling on the mappable genome alone and then impute copy number values on the low mappability regions using adjacent copy number calls.

The depth of coverage per 20kb mappable bin varies from 2 to 4 read pairs when calling events on a single cell, to as many as 2000-4000 read pairs when calling events on a cluster of 1000 similar cells. This makes necessary an algorithm that adapts to a large range of read depth. We define a depth-200 window $D_{200}$ per cell, roughly as the number of mappable bins that would have to be aggregated to reach a depth of 200 read pairs per window: $D_{200}$ = 200/Median($\{x_{\tilde{\alpha}} \mid x_{\tilde{\alpha}} > 0\}$). At this scale the relative error due to finite-depth random sampling effects is $1/200^{0.5} \approx 7\%$. The heuristics we employ below perform computations at this scale to minimize the impact of sampling effects.

***Copy number determination – GC correction:*** The coverage per bin is influenced by the GC content of the sequence. This effect arises from the library preparation reaction and the sequencer itself. Based on observing numerous samples we have found that the GC variation is best modeled as a quadratic function $f(g_\alpha;l,q) = 1 + l(g_\alpha - o) + q(g_\alpha - o)^2$, where $o$ = 0.45. We chose $o$ to be a conveniently chosen origin point with respect to which we normalize bins with lower and higher GC content, and $l,q$ are linear and quadratic coefficients to be determined. We estimate $l$ and $q$ on a per-cell basis. We aggregate reads from $D_{200}$ mappable bins to create a vector $y_\mu = \Sigma_{\mu D_{200} \le \tilde{\alpha} < (\mu+1) D_{200}} x_{\tilde{\alpha}}$, and then divide by the mean $z_\mu = y_\mu / \langle y_\mu \rangle$. We similarly computed the average GC content for bins of size $D_{200}$ as. For given values of $l$ and $q$, we calculate the normalized read counts $w_\mu(l, q) = z_\mu / f(g_\mu; l, q)$. Next, we calculated the normalized histogram $h(w\mu)$ and the entropy $E(w_\mu, l, q) = -\sum -h \log h$. To find the optimal l and q we minimize the objective function $O(l,q) = E(w_\mu, l, q) + \frac{1}{2}\lambda(l^2 + q^2)$. Here $\lambda$ = 0.005 is a regularization parameter that prevents solutions from a run-off to $\infty$.

**Copy number determination – identify candidate breakpoints:** We assume that the fluctuations of the copy number occur over a length scale larger than $D_{200}$. In other words, the genome is organized into segments of uniform copy number and there is a discontinuity at the junction of two such segments. We refer to these discontinuities as breakpoints. Our goal here is to identify an initial set of breakpoint locations, and we employ a log-likelihood ratio statistic to do so.

At each mappable bin $\tilde{\alpha}$, we define the left and right neighborhoods $N_L = [\tilde{\alpha} - D_{200}, \tilde{\alpha})$ and $N_R = [\tilde{\alpha}, \tilde{\alpha} + D_{200})$. The log-likelihood ratio (LLR) statistic is then defined at bin $\tilde{\alpha}$ as:

$$\mathrm{LLR}_{\tilde{\alpha}}(N_L, N_R) = \sum_{\tilde{\beta} \in N_L} \log \frac{\mathrm{Prob}(x_{\tilde{\beta}} \mid \mu_L)}{\mathrm{Prob}(x_{\tilde{\beta}} \mid \mu_{L \cup R})} + \sum_{\tilde{\beta} \in N_R} log \frac{\mathrm{Prob}(x_{\tilde{\beta}} \mid \mu_R)}{\mathrm{Prob}(x_{\tilde{\beta}} \mid \mu_{L \cup R})}$$

where $\mu_{NL}$, $\mu_{NR}$, $\mu_{NL \cup NR}$ are emission rates defined as:

$$\mu_{N_L} = \frac{\sum_{\tilde{\beta}\in N_L} x_{\tilde{\beta}}}{\sum_{\tilde{\beta}\in N_L} f(g_{\tilde{\beta}})}$$

$$\mu_{N_R} = \frac{\sum_{\tilde{\beta}\in N_R} x_{\tilde{\beta}}}{\sum_{\tilde{\beta}\in N_R} f(g_{\tilde{\beta}})}$$

$$\mu_{N_L \cup N_R} = \frac{\sum_{\tilde{\beta}\in N_L \cup N_R} x_{\tilde{\beta}}}{\sum_{\tilde{\beta}\in N_L \cup N_R} f(g_{\tilde{\beta}})}$$

and $\mathrm{Prob}(x \mid \mu) = \exp\left(x \log \mu - \mu - \log \Gamma\,(x+1)\right)$ is the probability mass function for a Poisson distribution.

Note that we circularized the genome so these neighborhoods can be consistently defined at the genome boundaries. When the bin $\tilde{\alpha}$ and its neighborhood $N_L \cup N_R$ have the same copy number $LLR_{\tilde{\alpha}}$ is close to 0 since the left and right neighborhoods are indistinguishable in terms of their count statistics. However, when $\tilde{\alpha}$ is near a breakpoint the statistic is positive. We choose a significance threshold of 5, which allows for a good balance of sensitivity and false positives in breakpoint detection **(Figure S1C)**. The initial set of breakpoints $B_0$ are thus defined as the set of local maxima of LLR that are above the significance threshold: $B_0 = \{\tilde{\alpha} \mid LLR_{\tilde{\alpha}} > LLR_{\tilde{\alpha}-1},\ LLR_{\tilde{\alpha}} < LLR_{\tilde{\alpha}+1},\ LLR_{\tilde{\alpha}} > 5\}$.

***Copy number determination – breakpoint refinement:*** Using every local maximum of the LLR statistic leads to many false positive breakpoints due to random fluctuations. We filter breakpoints by recomputing the LLR statistic as follows: assuming the elements of $B_0$ are sorted, for $b_n \in B_0$, we define left and right neighborhoods using the adjacent breakpoints $b_{n-1}, b_{n+1} \in B_0$ as $N_L = [b_{n-1}, b_n)$ and $N_R = [b_n, b_{n+1})$ and recompute the LLR statistic for each $b_n \in B_0$. We delete the breakpoint $b_n$ if the statistic $LLR_{bn} < 5$. We iterate through the list of candidates and delete breakpoints until every breakpoint $b$ satisfies $LLR_b >= 5$. This procedure gives us a smaller set of breakpoints $B_1$.

The initial set of breakpoints was identified using a fixed window size of $D_{200}$. This procedure in some cases can miss events smaller than $D_{200}$ that are in principle detectable. Furthermore, the breakpoint deletion procedure above while eliminating a large fraction of false positives also deletes true positive breakpoints. As a next step we scan the genome for evidence of a breakpoint as follows: we pick adjacent breakpoints $b_n, b_{n+1} \in B_1$ and compute the LLR statistic for every bin $b \in (b_n, b_{n+1})$ using the left and right neighborhoods $N_L = [b_n, b)$ and $N_R = [b, b_{n+1})$. If there is a bin $b$ with $LLR_b > 5$ we add the breakpoint $b$ to $B_1$. We iterate through the set of breakpoints $B_1$ and add breakpoints until there are no more breakpoints to be found. This defines a set of breakpoints $B_2$.

After generating an exhaustive list of breakpoints, we test whether each breakpoint can be moved by ±2 bins to improve the LLR statistic. This improves the accuracy of breakpoint location and the sensitivity of small event (< 200 kb) detection. We pick a triple of adjacent breakpoints $b_{n-1},b_n,b_{n+1} \in B_2$ and test whether $b_n$ can be replaced by a better location in the range $b \in [b_{n-2},b_{n+2}]$. A better location is defined as $b_n \rightarrow \underset{b\in[b_{n-2},b_{n+2}]}{\operatorname{argmax}} (\{LLR_b(N_L = [b_{n-1}, b), NR = [b, b_{n+1})\})$. We only perform this wiggle if $b_{n+1}-b_n > 2$ and $b_n-b_{n-1} > 2$. This gives us the final set of breakpoints $B$.

***Copy number determination – segment determination:*** The breakpoint set $B = \{b_1 = 0,\ldots,b_{|B|} = L_{GM} -1\}$ captures the set of bins where there is a copy number discontinuity. We add the origin 0 and the last genome bin $L_{GM} -1$ as mandatory breakpoints. We define segments as regions of constant copy number between adjacent breakpoints. The set of segments $S$ is defined as $\mathrm{S} = \{[\mathrm{b_n}, \mathrm{b}_{n+1}) \mid \mathrm{n} \in [1, |\mathrm{B}|)\}$. The segments in $S$ are a set of non-overlapping intervals that completely cover the genome.

***Copy number determination – fit to integer copy number:*** We associated each segment $s_a \in S$ with the length of the segment $l_a$ and the mean read count per $D_{200}$ window, $\mu_a$. Our goal here is to estimate the scale factor $\sigma$ for the cell, which in turn allows us to estimate the copy number $p_a$ for each segment. As we modeled $x_\alpha$ as a Poisson distribution, $x_\alpha$ is defined up to the transformation $p_a \rightarrow np_a$ and $\sigma \rightarrow \sigma/n$, where $n$ is an arbitrary integer. This transformation implies that there exists a fundamental solution such that all other solutions can be generated by integer multiplication. We describe here an algorithm to find the fundamental solution. In principle, the scale factor per cell σ could be inferred from the efficiency of the barcoding reaction and the sequencing depth, and defer this line of investigation to future work.

The scale can be estimated by minimizing the objective function $\mathrm{O}(\sigma) = \frac{1}{\mathrm{L_{GM}}} \sum_{\mathrm{a}\in[1,|\mathrm{S}|]} \mathrm{l_a} \sin^2\left(\frac{\pi\mu_\mathrm{a}}{\sigma}\right)$, which is minimized when each term is zero. This happens when $\mu_a = \sigma p_a$, $p_a \in \mathbb{Z}$. The function has multiple local minima and each one defines a copy number solution. Let $\sigma$ denote a local minimum of the objective function. The copy number of each segment $s_a \in S$ (and therefore for every 20 kb bin in $s_a$) is $\mathrm{n_a} = \mathrm{Round}\left(\frac{\mu_\mathrm{a}}{\sigma}\right)$, where Round is the integer rounding function. We can compute an average ploidy for each local minimum as $\sum_\mathrm{a} \mathrm{n_a l_a} / \mathrm{L_{GM}}$ and a copy number per bin as $\mathrm{p}_\alpha = \mathrm{n_a}, \alpha \in \mathrm{S_a}$.

Next, we filter away local minima that do not produce good integer fits to the data. To do so, we evaluate the quality of each candidate solution by computing a copy number-normalized noise measure. Let $\mathrm{P_i} = \{\tilde{\alpha} \in [0, \mathrm{L_{GM}} - 1] \mid \mathrm{p}_{\tilde{\alpha}} = \mathrm{i}\}$ denote the set of 20 kb bins with copy number $i = 1,2,\ldots$. For each copy number $i$, we partition the 20 kb bins in $P_i$ into $|P_i|/D_{200}$ bins of size $D_{200}$ and sum up the read counts in each partition. This creates a vector of read counts per $D_{200}$ window denoted by $c_i$. We expect these counts to be minimally dispersed around the mean beyond the amount dictated by Poisson statistics. We quantify by computing a depth-adjusted coefficient of variation for each vector $c_i$ as $\mathrm{DCV_i} = \sqrt{\left(\frac{\mathrm{Var(c_i)}}{\mathrm{Mean(c_i)^2}} - \frac{1}{\mathrm{Mean(c_i)}}\right)}$, which is then averaged over all copy number levels as $\mathrm{DCV} = \frac{\sum_i |P_i| \mathrm{DCV_i}}{\sum_i |P_i|}$. We discard solutions with high values of *DCV* as described below. Finally, we pick the global minimum of the remaining solutions as defining the copy number of the cell.

The global minimum of the objective function corresponds to the fundamental solution. Considering a *p*-ploid cell where the entire genome in a cell consists of one segment with copy number $p > 1$, and that the true scale factor for this cell is $\sigma_0$, the objective function then consists of a single term $O = \sin^2(\pi\mu 1/\sigma)$. The variable $\mu_1$ is normally distributed when $\mathrm{L_{GM}} \gg 1$ by the central limit theorem with $E(\mu_1) = p\sigma_0$ and $Var(\mu_1) = p\sigma_0/L_{GM}$. The objective function then can be related to the characteristic function of the normal distribution and $\langle \mathrm{O}(\sigma) \rangle = \frac{1}{2}\left(1 - cos\frac{2\pi p\sigma_0}{\sigma} \exp\left(-\frac{2\pi^2 p\sigma_0}{\mathrm{L_{GM}}\sigma^2}\right)\right)$. Comparing the values of the objective function $O(\sigma = \sigma_0)$ (the correct answer) and at $O(\sigma = p\sigma_0)$ we see that the latter has a smaller value. In other words, the true solution has been disfavored relative to a lower integer multiple. This example serves to illustrate why the objective function in general picks the fundamental solution. However, the absolute minimum value of the objective function does not always produce the correct answer. For example, in a perfectly diploid female human cell where all the chromosomes are at copy number 2, the algorithm would put all the chromosomes at copy number 1, which is obviously not correct. We handle this as a special case.

***Copy number determination – imputation on non-mappable regions:*** Finally, we impute the copy number values for low mappability bins using neighboring mappable bins. Let *R* denote a maximal contiguous region of low mappability. By definition the bins on the left and right of *R* are mappable and their copy numbers have been estimated by the algorithm described above. When the left and right bin have identical copy numbers *p* and when the size of *R* is less than 500 kb we assign the bins in *R* a copy number of *p*. When the size of *R* is 500 kb or above, or when the left and right bins have unequal copy numbers we do not make a copy number call over *R*.

**SUPPLEMENTARY METHODS – Cell cycle analysis**

***Assigning cell cycle state to scDNA-sequenced cells:*** For a given sample, we classified the genome of each sequenced cell $i \in I$ to one of three states (G0/G1, S, apoptotic) as follows. Under the assumption that the G0/G1 population is larger than any of the other populations, we defined the G0/G1 ploidy, $p_{g0g1}$, as the median ploidy across all sequenced cells of a given sample. We then calculate three features for each cell, $x$: i) its distance, $d_x$, to $p_{g0g1}$; ii) its total number of breakpoints, $b_x$, and iii) the Pearson correlation coefficient, $r_x$, between the number of rare breakpoints observed in the cell per each chromosome and the number of replication origins per chromosome. Rare breakpoints were defined as breakpoints that were shared among less than 1% of cells.

We distinguished G0/G1 cells from cells with higher genome fragmentation. We divided cells into two groups – $P_A := \{x \in I \mid p_x \geq p_{g0g1}\}$ and $P_B := \{x \in I \mid p_x < p_{g0g1}\}$ – containing cells above and below the sample's G0/G1 ploidy respectively. We fitted two sigmoid functions, one for each subgroup, to model cell ploidy as function of the number of breakpoints per cell: $p_i \sim \begin{cases} f_A(b_i), if\ i \in P_A \\ f_B(b_i), if\ i \in P_B \end{cases}$

We then calculated $B \coloneqq argmin_b | f_A(b) - f_B(b)|$, as the threshold distinguishing G0/G1 cells from apoptotic cells:

$$apoptotic := \{x \in P_B \mid b_x \geq B\}$$

and from replicating cells:

$$S := \{x \in P_A \mid b_x \geq B\}$$

I.e. the yet unclassified cells were assigned to the G0/G1 state:

$$G0G1 := \{x \in I \mid b_x < B\} - \{S \cup apoptotic\}$$

The highest correlation to replication origins was observed for replicating cells (***SI Appendix, Fig. S*5B-J**), supporting the accuracy of above cell cycle phase assignment strategy. We removed cell cycle specific breakpoints from further analysis, keeping only those breakpoints present among at least 1% of G0/G1 cells and encompassing segments of at least 5 Mb. Population-average copy number per segment per sample was calculated as the mean copy number across G0/G1 cells of that sample.

***Inferring clonal dynamics from distribution of replicating cells among clones:*** For each detected and confirmed clone of a given sample, we calculated whether its % replicating cell assignment was different than expected by chance from its G0/G1 representation. Hereby we excluded clones below 4% size, because their absolute cell count was too small to reliably perform these calculations. To infer positive selection, we used the hypergeometric distribution and calculated the p-value of sampling at least the observed number of replicating clone members, ***q***, as:

P = phyper(***q, m, n, k***)

where $k$ is the total number of cells sampled from the clone, $n$ is the number of clone members in G0/G1 and $m$ is the expected number of clone members that are replicating (assuming proportionality to G0/G1 clone size). The p-value of sampling maximum $q$ replicating cells was calculated by subtracting above value from 1, and was used to infer negative selection. P-values were adjusted for multiple hypotheses testing using the FDR method (R function "p.adjust").

**SUPPLEMENTARY METHODS – single cell RNA-Seq**

***ScRNA-Seq library preparation and sequencing:*** We used the Chromium Controller instrument (10X Genomics Inc., Pleasanton, CA) and the Single Cell 3' Reagent kit (v2) to prepare individually barcoded single cell RNA-Seq libraries following the manufacturer's standard protocol. Briefly, single cell suspensions were loaded on a Chromium Controller instrument and were partitioned in droplets. Reverse transcription is performed, followed by droplet breaking, and cDNA amplification. Each cDNA molecule thus contained the read 1 sequencing primer, a 16bp cell-identifying barcode, and a 10bp UMI sequence [20]. We performed enzymatic fragmentation, end-repair, and a-tailing followed by ligation of a single-end adapter containing the read 2 priming site. PCR was performed using the Illumina P5 sequence and a sample barcode as described earlier. Libraries were purified with SPRIselect beads (Beckman Coulter, Brea, CA) and size-selected to ~450bp. Finally, sequencing libraries were quantified by qPCR before sequencing on the Illumina platform using 26x98 paired-end reads. The Cellranger software suite was used to process scRNA data, sample demultiplexing, barcode processing, and single cell 3' gene counting. The cDNA insert, which is contained in the read 2, was aligned to the GRCh38 human reference genome. Cellranger provided a gene-by-cell matrix, which contains the read count distribution of each gene for each cell.

***ScRNA-Seq data preprocessing:*** We used a curated set of seven biological and technical features to detect and remove low-quality cells [20, 34]. Biological features included: 1) transcriptome variance and expression of 2) cytoplasm localized genes, 3) mitochondrially localized genes, 4) mtDNA encoded genes. Technical features included: 5) % mapped reads, 6) % multi-mapped reads and 7) %non-exonic reads (intergenic & intronic). These features robustly identify low quality cells independently of cell type and of the experimental setting. The analysis was performed using "Celloline" [34] and the R-package "Cellity" [34]. For additional processing, we used the software suite Seurat (v2.3.2) [35]. Briefly, UMI counts were capped at the 99% quantile and only cells expressing at least 1,000 genes were included in subsequent analysis. Also, most cells classified by Cellity as low quality had a high percentage expressed mitochondrial genes as quantified by Seurat (***SI Appendix, Fig. S*6**).

***Assigning cell cycle state to scRNA-sequenced cells:*** Leveraging prior knowledge in form of cell-cycle annotated genes and deploying a rank-based comparison across single cells, has been shown to robustly capture the transcriptional cell-cycle signature across different cell types and experimental protocols [23]. We employed such pathway-centric approach to classify the transcriptome of each sequenced cell to a cell cycle state as follows:

***Pathway quantification:*** The gene membership of 1,417 pathways was downloaded from the Reactome database [36] (v63). First the transcriptome profiles of high quality cells detected within a given sample were scaled to the number of UMIs per cell (Seurat function "ScaleData"). We used the GSVA function [37] to model variation in pathway activity across cells of the sample (R function "gsva", mx.diff=TRUE). GSVA starts by evaluating the expression magnitude of a given gene in a given cell, in the context of the sample population distribution. To reduce gene specific biases (i.e. caused by GC content and gene length), an expression-level statistic was calculated for each gene from a kernel estimation of its cumulative density function. GSVA then calculated a rank-based, cell specific enrichment scores using the Kolmogorov-Smirnov like random walk statistic. For any given sample, pathways for which less than ten gene members were expressed in the scRNA-Seq data were not quantified.

***Quantification of cell cycle pathways activity:*** The gene membership of 39 cell cycle pathways was downloaded from the Reactome database [36] (v63), whereby each pathway consisted of at least ten genes **(*SI Appendix,* Table S4)**. We used the GSVA method [37] to model variation in pathway activity across cells of a given cell line, as described later.

***Pathway and cell classification***: Pathways were classified into three groups depending on their main activation timing during: i) G0/G1 (10 pathways, further referred to as $P_{G0G1}$); ii) S (5 pathways, further referred to as $P_S$) and iii) G2M (26 pathways, further referred to as $P_{G2M}$). Each class was normalized by its maximum activity across cells. As previously described, the 39 pathways were used as features to perform hierarchical clustering of cells (Euclidean distance metric and ward.D2 agglomeration method) into four clusters C:={$C_1$, $C_2$, $C_3$, $C_4$}. To classify each cluster $x \in C$ as either an G0/G1, S or G2M representative, we tested 39 null-hypotheses, one for each pathway p, namely that the activity of p in cells from x exceeds the activity of p in cells from {C – x}. We tested our hypotheses using the

Wilcox rank-sum test and p-values were adjusted for multiple testing. For each pathway class $\delta \in \{G0G1, S, G2M\}$ we calculated the average effect size as:

$$P(x|\delta) := \frac{1}{|P_\delta|} \sum_{p \in P_\delta} e_{p,x}, \text{ where:}$$

$$e_{p,x} = \begin{cases} effect\ size, if\ Wilcox\ p \leq 0.05 \\ 0, otherwise \end{cases}$$

Finally, we assigned cell cycle phase $argmax_{\delta \in \{G0G1,S,G2M\}} P(x|\delta)$ to each cluster $x \in C$.

***Assigning scRNA-sequenced single cells to cell types in P5931:*** We employed the method of Macosko *et al.* [40] to rank genes based on their normalized dispersion (Seurat [35, 41] function *FindVariableGenes*). We used 1,097 highly-variable genes to compute 60 principal components (Seurat function RunPCA). The first 51 principal components explained >90% of the variance in the data and were used as input for the subsequent tSNE analysis (Seurat function RunTSNE).

**SUPPLEMENTARY METHODS – LIAYSON: Calling CNVs from scRNA-Seq**

To infer a cell's copy number state at any given locus, the LIAYSON algorithm uses a cell's read counts across the entire genome, thereby mitigating the influence of non-genetic factors on mRNA expression. In addition to raw UMI counts, LIAYSON requires as input the population-average (bulk) segmentation profile of the sample and the classification of cells into G0/G1, S and G2M subsets. It consists of two steps – aggregating expression across copy number segments, and calling copy number from segmental expression. These two steps are detailed as follows.

*Calculating cell-by-segment expression matrix.* Let $S := \{S_1, S_2, \ldots S_n\}$ be the set of $n$ genomic segments that have been obtained from DNA-sequencing $i \in I$ cells of given sample (e.g. from bulk exome-sequencing, scDNA-sequencing, etc.). To prepare each cell's RNA-seq profile for copy number analysis we first grouped genes by their segment membership, such that $\boldsymbol{E_{ij}}$ and $\boldsymbol{G_{ij}}$ are the average number of UMIs and the number of expressed genes per segment $S_j$ per cell *i*.

To reduce data sparsity and the effect of non-genetic factors on gene expression we excluded genomic segments shorter than 10 Mb. For each cell cycle phase (**G0/G1, S, G2/M**), we also excluded genomic segments j for which $\overline{\boldsymbol{G_{*j}}}$ – the average number of expressed genes per cell – was below 20 (setting this threshold too high would also exclude single copy losses).

*Calling copy numbers from G0/G1 cell-by-segment expression matrix.* We first normalize the cell-by-segment expression matrix to gene coverage, by fitting a linear regression model for each j ∈ S:

$E_{*j} \sim Z_{*}$, where $Z_i \coloneqq \sum_{j \in S} G_{ij}$ – is the overall gene coverage of a given cell.

The model's residuals $R_{ij}$ reflect inter-cell differences in expression per segment that cannot be explained by differential gene coverage per cell. A first approximation of the cell-by-segment copy number matrix C is then given by: $\mathbf{C_{ij}}$ := $R_{ij}$ * ($cn_j$ / $\mu_j$ ), where $\mu_j \coloneqq \frac{1}{|I|} \sum_{i \in I} R_{ij}$, is the mean residual per segment across cells and $cn_j$ is the G0/G1 population-average copy number of segment j derived from DNA-seq. Above transformation of $\mathbf{E_{ij}}$ into $\mathbf{C_{ij}}$ is in essence a numerical optimization, shifting the distribution of each segment to the average value expected from bulk DNA sequencing.

Let x' ∈ C be the measured copy number of a given cell-segment pair, and x its corresponding true copy number state. The probability of assigning copy number x to a cell i at locus j depends on:

A. **Cell i's read count at locus j**, calculated conditional on the measurement x'.
   We fit a Gaussian kernel on the read counts at locus j across cells to identify the major (M) and the minor (m) copy number states of j as the highest and second highest peak of the fit respectively. Then we calculate the proportion of cells expected at state m as: $f \coloneqq \frac{cn_j - M}{m - M}$. The probability of assigning copy number x to a cell i at locus j is calculated as:

$$\mathrm{P_A}(x|x') \coloneqq \begin{cases} 0, if\ x \notin \{m, M\} \\ \mathrm{P_{ij}}(\,\mathrm{x}'|\mathrm{N}(m, sd = f\,)\,), if\ x == m \\ \mathrm{P_{ij}}(\,\mathrm{x}'|\mathrm{N}(M, sd = 1 - f\,)\,), if\ x == M \end{cases}$$

B. **Cell i's read count at other loci**, i.e. how similar the cell is to other cells that have copy number x at locus j. We use Apriori – an algorithm for association rule mining – to find groups of loci that tend to have correlated copy number states across cells. Let $R^j_{K \to x}$ be the set of rules concluding copy number x for locus j, where k ∈ K are copy number profiles of up to n=4 loci in the form {$S_1$=$x_1$, $S_2$=$x_2$, … $S_n$=$x_n$}. For each cell $i \in I$ corresponding to any of the copy number profiles in K, we calculate:

$$\mathrm{P_B(x)} \sim \sum_{r \in \mathrm{R}^j_{K \to x}} C_r$$, the cumulative confidence of the rules in support of x at j.

We first obtain a seed of cell-segment pairs by assigning a-priori copy number states only when $\mathrm{argmax}_{x \in [1,8]}\ \mathrm{P_A}(x|x'') > t$. We use this seed as input to B. Finally, a-posteriori copy number for segment j in cell i is calculated as: $argmax_{x \in [1,8]}\ P_A(x|x'') + P_B(x)$.

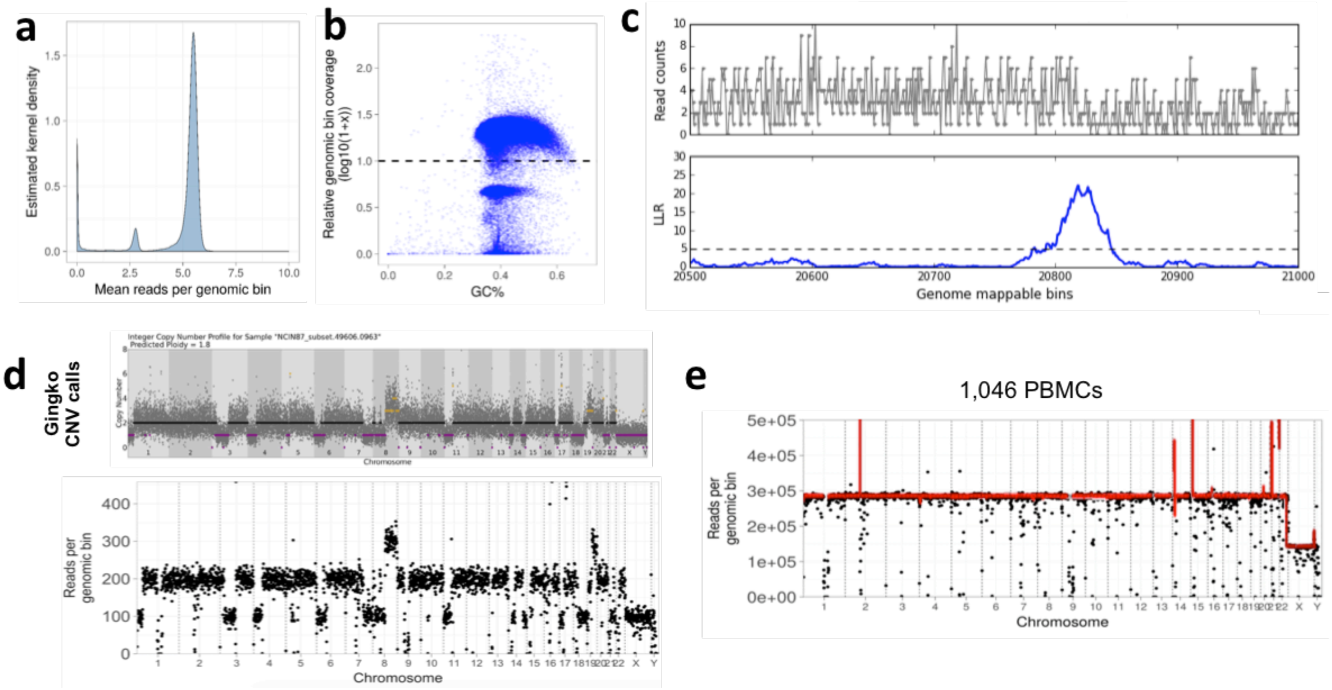


**Figure S1: Technical performance of scDNA-Seq.** (**a**) Reads per genomic bin. For each 20kb bin in GRCh38, the average number of reads across 1,046 cells was calculated. (**b**) GC bias in scDNA-Seq coverage. The relative coverage in each 20kb bin in GRCh38 was normalized by the average coverage per bin for each cell, and then averaged across 1,046 PBMCs. Dashed line: Relative coverage of 1, reflecting average per-bin coverage. (**c**) Candidate copy number breakpoint detection. Discontinuities in read counts is detected by the log-likelihood ratio (LLR) statistic across genome-wide bins. A candidate breakpoint is identified by a peak in the LLR statistic. (**d**) Copy number landscape of NCI-N87 cell 963 determined by Ginkgo (top) is shown alongside binned read coverage (bottom). (**e**) Copy number calls and average read coverage for 1,046 PBMCs. Each point indicates the binned read coverage, while the red line indicates the average copy number call for that region across cells.

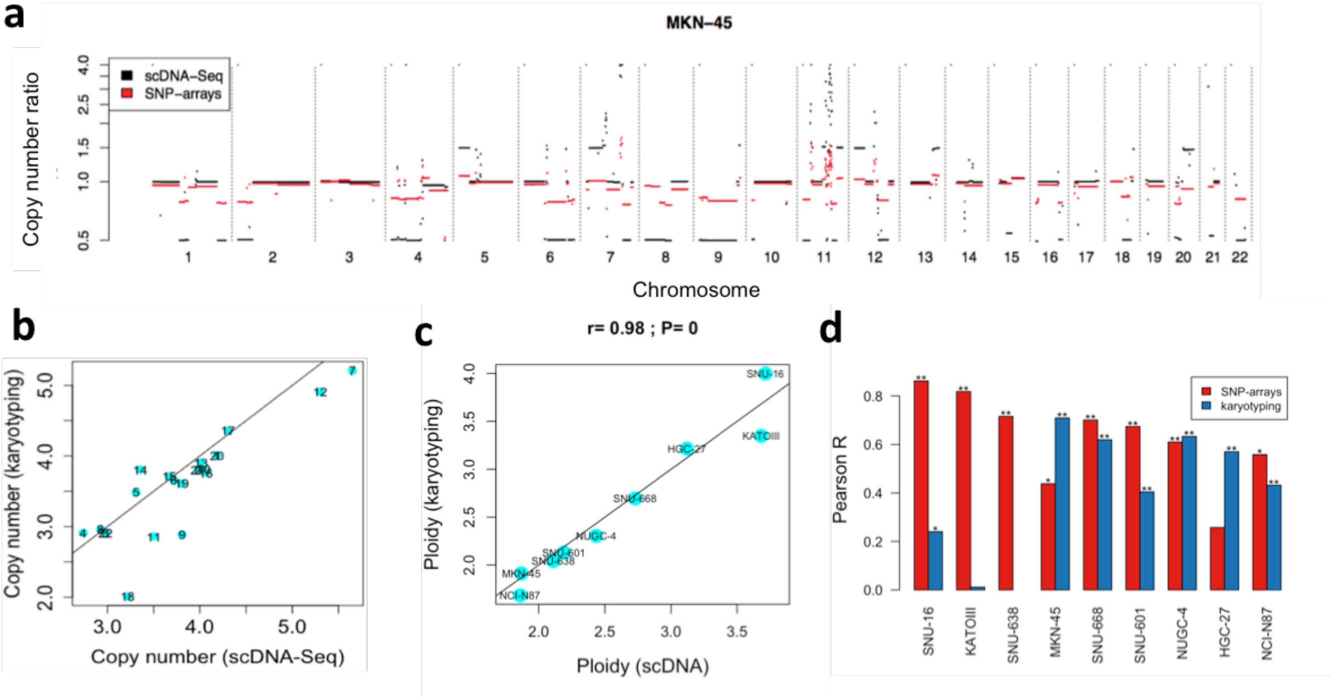


**Figure S2: Karyotyping and SNP-array analysis confirm scDNA-Seq derived aneuploidy.** (**a**) ScDNA-Seq derived aneuploidy of MKN-45 (black) is confirmed by SNP-array data (red). (**b**) Correlation between average scDNA-Seq derived (x-axis) and karyotyping derived (y-axis) copy number per chromosome is shown for SNU-16 (Pearson r=0.86; P<1E-5). (**c**) Average ploidy per cell line inferred by cellranger-dna from scDNA-Seq (x-axis) correlates to cell line's Karyotype (y-axis) (Pearson r=0.98; P<1E-5). (**d**) ScDNA-Seq derived copy number per segment correlates with karyotyping and with SNP-arrays, as is shown for MKN-45 and SNU-16 along with the other cell lines (**: P<=0.005; *: P<=0.05). In general, SNP-array data was obtained from older passages of the respective cell lines and therefore had weaker correlation to the scDNA-Seq data than did karyotyping. SNP-array data was not available for SNU-638.

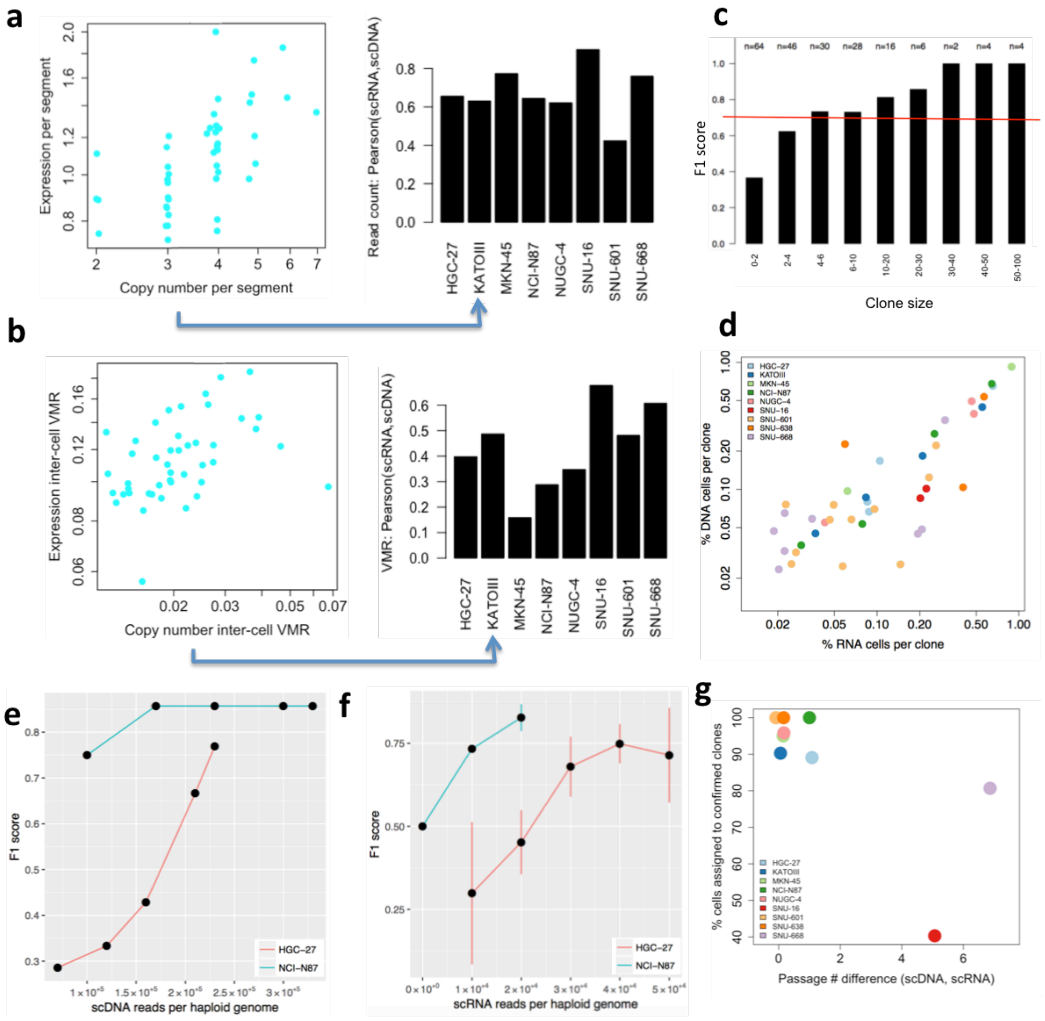


**Figure S3: ScDNA- and RNA-Seq mutual validation.** (**a-b**) **Validation at meta-population resolution**. (**a**) Average expression per segment across KATOIII cells (left) reflects the cell population's average copy number (Pearson r=0.63; P=8.6E-6). Correlation coefficients between scDNA- and scRNA-Seq derived average read counts per segment are shown for KATOIII along with the other seven gastric cancer cell lines (right). (**b**) Variance-to-mean ratio (VMR) of expression per segment across KATOIII cells (left) correlates with the cells' variability in copy number states (Pearson r=0.49; P=8.0E-4). Correlation coefficients between scDNA- and scRNA-Seq derived VMR are shown for all gastric cancer cell lines (right). (**c-f**) **Validation at subpopulation resolution**. (**c**) Clone detection F1 score increases with increasing clone size across all nine cell lines. (**d**) The relative proportions of scDNA-Seq (y-axis) and scRNA-Seq (x-axis) cells per clone correlate within and across cell lines. (**e-f**) Dependence of clone detection F1 score on sequencing depth in NCI-N87 and HGC-27. F1 score increases with increasing number of reads sequenced per each cell's haploid genome. F1 score of scDNA-Seq (**e**) and scRNA-Seq (**f**) clone detection was calculated using the respectively other technique as control. For (**e,f**), between 20% and 95% of reads were sampled randomly for each of the two CLs and used to estimate performance at different sequencing depths. (**g**) Differences in passage number between scDNA- and scRNA-Seq experiments accompany differences in clonal composition observed between the two techniques for SNU-16 and SNU-668 (Pearson r=-0.71; P=0.032).

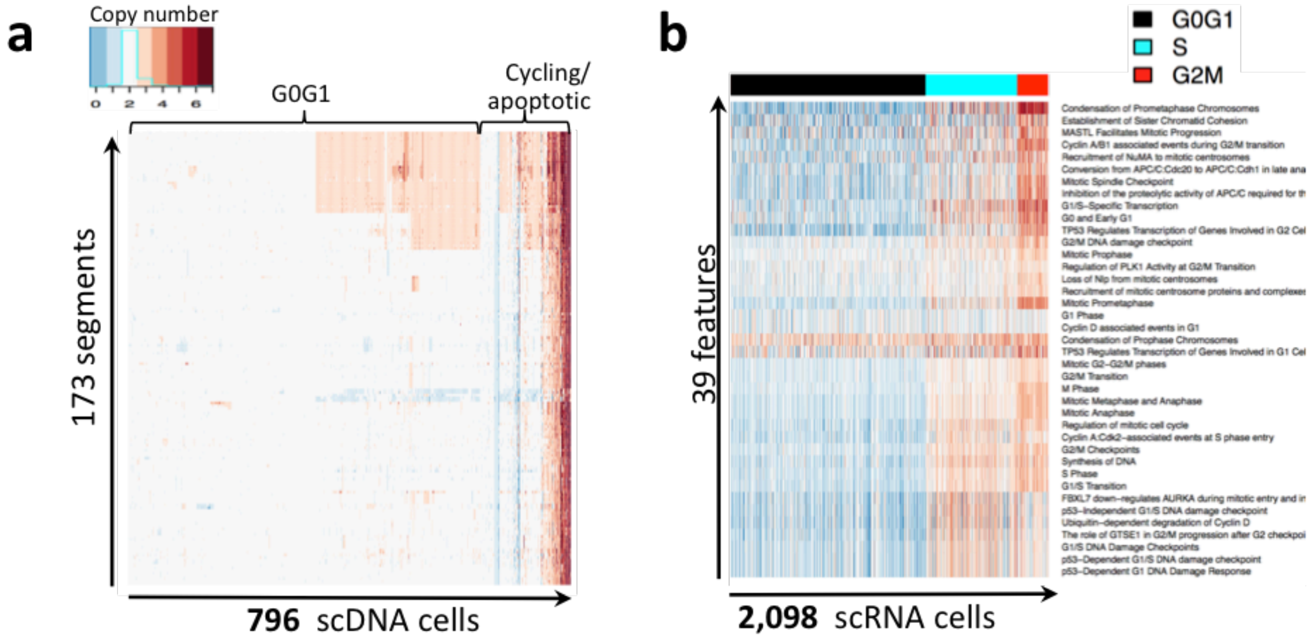


**Figure S4: Cell cycle phase assignment of sequenced cells isolated from a primary gastric cancer sample of patient P5931.** (**a**) Cell cycle assignment of 796 scDNA-sequenced cells (columns). (**b**) 2,098 scRNA-sequenced cells, isolated from the same primary gastric cancer sample, are clustered according to the activity of 39 pathways related to various states along the cell cycle (rows). Clusters are then classified as containing either G0/G1 cells (black), cells in S-phase (cyan) or cells in G2M (red).

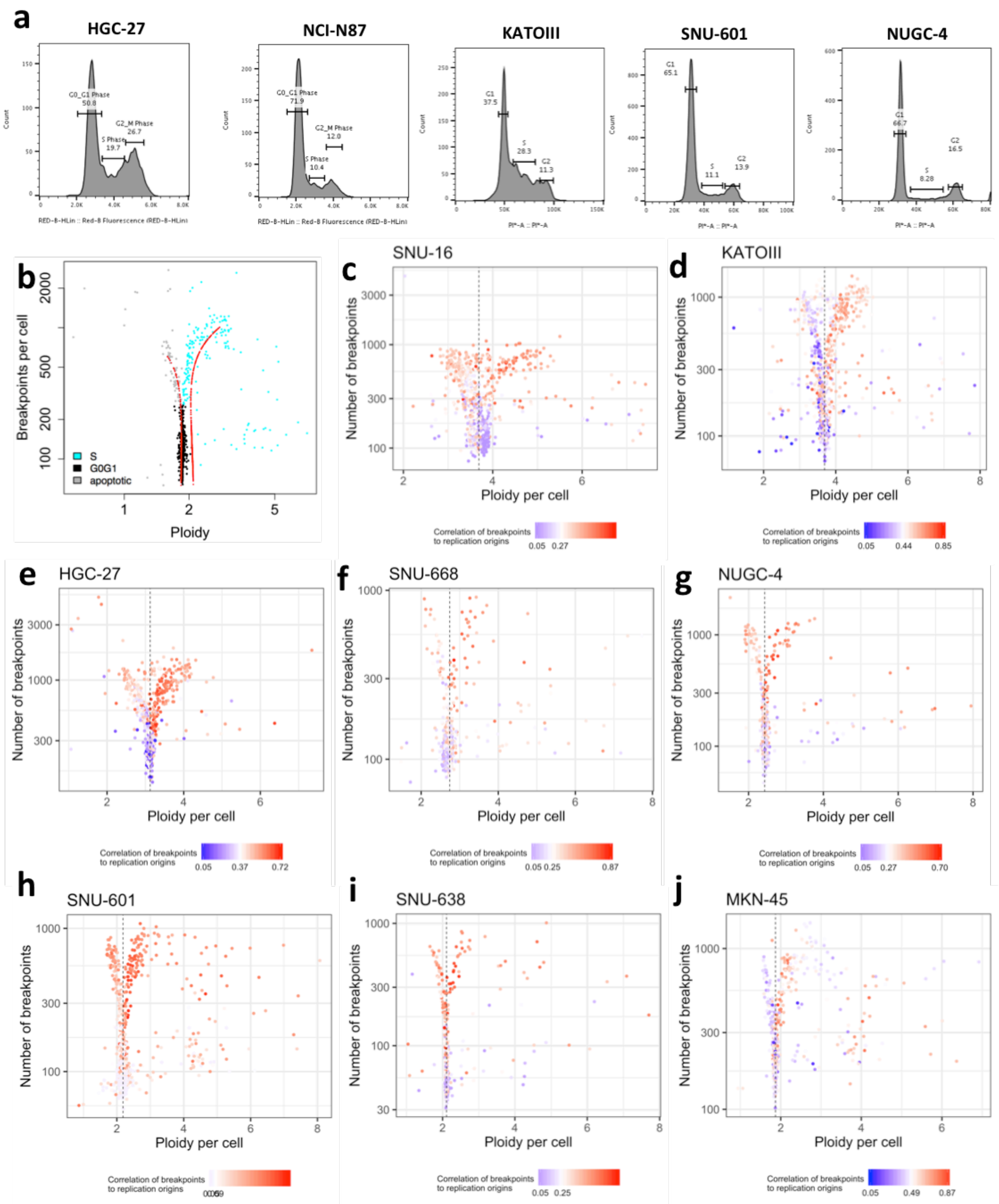


**Figure S5: ScDNA-Seq cell cycle phase assignment strategy and validation**. (**a**) Cell cycle analysis of five gastric cancer cell lines by flow cytometry. (**b**) Demonstration of cell cycle phase assignment strategy with scDNA-Seq using NCI-N87 as an example. Two sigmoid functions (dotted red lines) are fitted to model cell ploidy (x-axis) as a function of the number of breakpoints per cell (y-axis), for cells above- (right) and below (left) median sample ploidy. Cell cycle phase assignment according to the fitted functions **(Online Methods)** is color-coded. (**c-j**) Validation of assignment strategy using a third, independent feature: correlation to replication origins **(Online Methods)**. The highest correlation was observed for S cells, whereas G0/G1 cells had the lowest values as would be expected from cells that are not actively replicating.

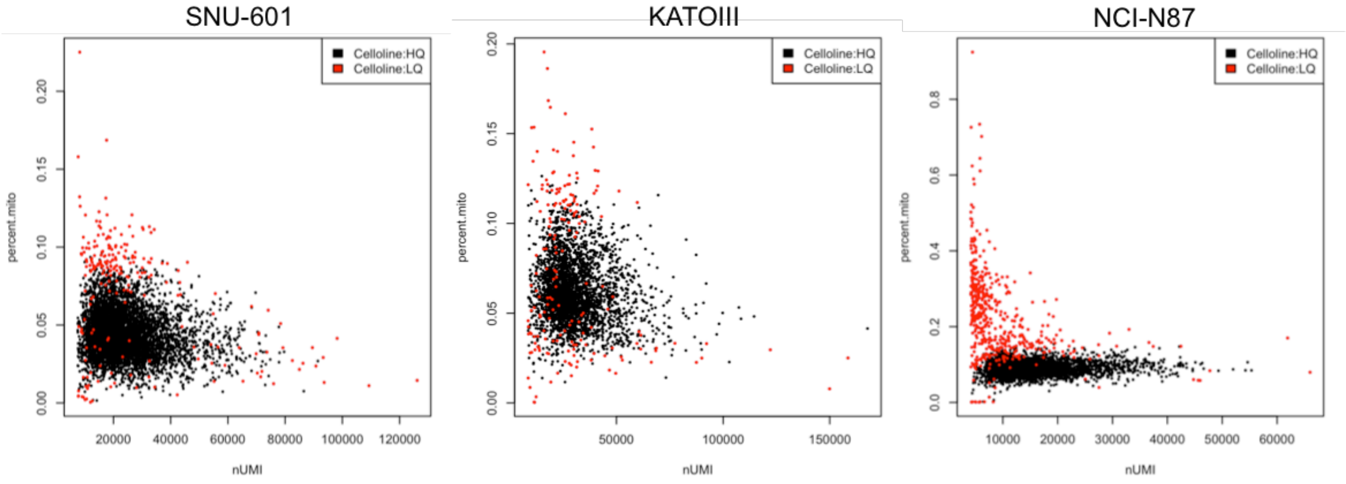


**Figure S6:** Cells excluded by Cellity due to low quality also have a high percentage expressed mitochondrial genes as quantified by Seurat, supporting the exclusion of these cells from further analysis.

**Table S1: ScDNA-Seq statistics**.

| | SNU-16 | KATOIII | HGC-27 | SNU-668 | NUGC-4 | SNU-601 | SNU-638 | MKN-45 | NCI-N87 | P5931 |
|---|---|---|---|---|---|---|---|---|---|---|
| **Sample type** | Cell line | Cell line | Cell line | Cell line | Cell line | Cell line | Cell line | Cell line | Cell line | Primary |
| **Sequenced cells** | 825 | 973 | 912 | 1,238 | 829 | 1,531 | 724 | 787 | 1,005 | 796 |
| **Mean reads per cell** | 1,006,726 | 1,213,316 | 1,458,391 | 585,062 | 1,478,651 | 711,598 | 639,428 | 916,841 | 1,321,999 | 892,532 |
| **Mean coverage per cell** | 0.06 | 0.07 | 0.09 | 0.04 | 0.09 | 0.04 | 0.04 | 0.06 | 0.08 | 0.05 |
| **Ploidy (scDNA-Seq)** | 3.71 | 3.68 | 3.12 | 2.73 | 2.43 | 2.19 | 2.11 | 1.87 | 1.86 | 2.19 |
| **Breakpoint count (all cells)** | 2,037 | 2,545 | 5,497 | 1,621 | 1,477 | 1,198 | 425 | 3,677 | 1,306 | 268 |
| **Breakpoint count (G0G1 cells)** | 96 | 86 | 144 | 92 | 61 | 81 | 35 | 118 | 60 | 28 |
| **G0G1 (%)** | 58% | 62% | 63% | 82% | 79% | 77% | 74% | 68% | 74% | 80% |
| **S (%)** | 17% | 21% | 18% | 10% | 11% | 14% | 12% | 22% | 18% | 14% |
| **Apoptotic (%)** | 25% | 17% | 19% | 8% | 10% | 9% | 14% | 10% | 8% | 7% |

**Table S2**: Passage number, confluence and general information about nine gastric cancer cell lines. Confluence was typically similar between scDNA- and scRNA-Seq experiments (80-90%), but diverged considerably for NUGC-4, explaining the discrepancy in the % cycling cells between the two techniques for this cell line (**Fig. 2b**).

| | | SNU-16 | KATOIII ❖ | HGC-27 | SNU-668 | NUGC-4 | SNU-601 ❖ | SNU-638 ❖ | MKN-45 | NCI-N87 |
|---|---|---|---|---|---|---|---|---|---|---|
| **Passage # \| % confluence** | **> Karyotyping** | P8 | P2 | P7(o) | P2 | P24(o) | P2 | P2 | P24(o) | P2 |
| | **> ScDNA-Seq** | P7 | P4 \| 65 | P7(o) \| NA | P9 \| 80-90 | P23(o) \| 20-30 | P3 \| 80 | P1 \| 50-60 | P23(o) \| NA | P3 \| NA |
| | **> ScRNA-Seq** | P2 | P4 \| 65 | P7(o) \| 80-90 | P2 \| 80-90 | P23(o) \| 80-90 | P3 \| 80 | P1 \| 50-60 | P23(o) \| 80-90 | P2 \| 80-90 |
| **Doubling time (h)** | | 27 | 32 | 17 | 74 | 36 | 47 | 58 | 24 | 47 |
| **Ploidy (Karyotyping)** | | 3.76 | 3.71 | 3.29 | 2.83 | 2.60 | 2.29 | 2.25 | 1.95 | 1.94 |
| **Age** | | 33 | 55 | NA | 68 | 35 | 34 | 48 | 62 | NA |
| **Sex** | | Female | Male | Unspecified | Male | Female | Male | Male | Female | Male |
| **Tumor Differentiation** | | Carcinoma | Carcinoma | Carcinoma | Carcinoma | Adenocarcinoma | Carcinoma | Carcinoma | Adenocarcinoma | Carcinoma |
| **Growth type** | | suspension | adherent + suspension | adherent | adherent | adherent | adherent | adherent | adherent + suspension | adherent |
| **Year of 1st report** | | 1990 | 1974 | 1976 | 1997 | 1988 | 1997 | 1997 | 1976 | 1990 |

(o) passage number includes that from vendor

❖ scDNA & scRNA sequenced from the same suspenion

**Table S3: ScRNA-Seq statistics**.

| | SNU-16 | KATOIII | HGC-27 | SNU-668 | NUGC-4 | SNU-601 | SNU-638 | MKN-45 | NCI-N87 | P5931 |
|---|---|---|---|---|---|---|---|---|---|---|
| **Sample type** | Cell line | Cell line | Cell line | Cell line | Cell line | Cell line | Cell line | Cell line | Cell line | Primary |
| **Replicates** | 2 | 1 | 2 | 1 | 1 | 1 | 1 | 1 | 1 | 1 |
| **Sequenced cells*** | 2,088 | 3,084 | 1,808 | 5,319 | 3,797 | 5,186 | 867 | 2,814 | 3,246 | 2,098 |
| **Mean reads per cell** | 138,104 | 76,707 | 150,155 | 51,949 | 53,046 | 84,420 | 71,714 | 54,031 | 40,555 | 130,035 |
| **Mean genes per cell** | 5,683 | 4,851 | 5,584 | 3,920 | 3,686 | 4,464 | 3,841 | 3,611 | 3,299 | 3,889 |
| **G0G1** | 28% | 49% | 29% | 72% | 47% | 64% | 79% | 55% | 72% | 61% |
| **G2M** | 37% | 18% | 21% | 9% | 25% | 13% | 9% | 12% | 11% | 10% |
| **S** | 35% | 33% | 51% | 19% | 28% | 24% | 11% | 33% | 17% | 29% |
| **Cycling** | 72% | 51% | 71% | 28% | 53% | 36% | 21% | 45% | 28% | 39% |

*Low quality cells excluded

**Table S4: Cell cycle pathways**. A total of 39 cell cycle pathways from the REACTOME database are listed along with their activation timing during S, G2M or G0/G1 phases of the cell cycle.

| | Pathway | Class |
|---|---|---|
| 1 | Establishment of Sister Chromatid Cohesion | S |
| 2 | Cyclin A:Cdk2-associated events at S phase entry | S |
| 3 | S Phase | S |
| 4 | Synthesis of DNA | S |
| 5 | Ubiquitin-dependent degradation of Cyclin D | S |
| 6 | Condensation of Prometaphase Chromosomes | G2M |
| 7 | Condensation of Prophase Chromosomes | G2M |
| 8 | Conversion from APC/C:Cdc20 to APC/C:Cdh1 in late anaphase | G2M |
| 9 | Cyclin A/B1 associated events during G2/M transition | G2M |
| 10 | FBXL7 down-regulates AURKA during mitotic entry and in early mitosis | G2M |
| 11 | G2/M Checkpoints | G2M |
| 12 | G2/M DNA damage checkpoint | G2M |
| 13 | G2/M Transition | G2M |
| 14 | Inhibition of the proteolytic activity of APC/C required for the onset of anaphase by mitotic spindle | G2M |
| 15 | Loss of Nlp from mitotic centrosomes | G2M |
| 16 | M Phase | G2M |
| 17 | MASTL Facilitates Mitotic Progression | G2M |
| 18 | Mitotic Anaphase | G2M |
| 19 | Mitotic G2-G2/M phases | G2M |
| 20 | Mitotic Metaphase and Anaphase | G2M |
| 21 | Mitotic Prometaphase | G2M |
| 22 | Mitotic Prophase | G2M |
| 23 | Mitotic Spindle Checkpoint | G2M |
| 24 | Recruitment of NuMA to mitotic centrosomes | G2M |
| 25 | Recruitment of mitotic centrosome proteins and complexes | G2M |
| 26 | Regulation of PLK1 Activity at G2/M Transition | G2M |
| 27 | Regulation of mitotic cell cycle | G2M |
| 28 | TP53 Regulates Transcription of Genes Involved in G2 Cell Cycle Arrest | G2M |
| 29 | The role of GTSE1 in G2/M progression after G2 checkpoint | G2M |
| 30 | Cyclin D associated events in G1 | G0G1 |
| 31 | G0 and Early G1 | G0G1 |
| 32 | G1 Phase | G0G1 |
| 33 | G1/S DNA Damage Checkpoints | G0G1 |
| 34 | G1/S Transition | G0G1 |
| 35 | G1/S-Specific Transcription | G0G1 |
| 36 | TP53 Regulates Transcription of Genes Involved in G1 Cell Cycle Arrest | G0G1 |
| 37 | p53-Dependent G1 DNA Damage Response | G0G1 |
| 38 | p53-Dependent G1/S DNA damage checkpoint | G0G1 |
| 39 | p53-Independent G1/S DNA damage checkpoint | G0G1 |

**Table S5: Mutual validation of scDNA- and scRNA-Seq derived clone identification.**

| | SNU-16 | KATOIII | HGC-27 | SNU-668 | NUGC-4 | SNU-601 | SNU-638 | MKN-45 | NCI-N87 | P5931 |
|---|---|---|---|---|---|---|---|---|---|---|
| Passage # difference (scDNA, scRNA) | 5 | 0 | 1 | 7 | 0 | 0 | 0 | 0 | 1 | NA |
| Clone # (scDNA) | 11 | 5 | 5 | 10 | 4 | 12 | 4 | 2 | 4 | 4 |
| Clone # (scRNA) | 7 | 5 | 5 | 10 | 4 | 11 | 3 | 3 | 4 | 4 |
| Confirmed clone membership (%) | 40.3 | 90.3 | 89.1 | 80.7 | 95.8 | 100.0 | 100.0 | 95.0 | 100.0 | 93.8 |
| TP # | 2 | 4 | 4 | 8 | 3 | 11 | 3 | 2 | 4 | 3 |
| FP # | 5 | 1 | 1 | 2 | 1 | 0 | 0 | 1 | 0 | 1 |
| FN # | 6 | 1 | 1 | 2 | 1 | 1 | 1 | 0 | 0 | 1 |

**Table S6**: Multiple regression model of the clone count per cell line as a function of ploidy and time in culture. Coefficients were calculated by fitting a least square linear regression on the nine gastric cancer cell lines. Number of clones per cell line increases with ploidy and decreases with the number of years since a cell line was first established.

| | Coefficient | Std. Error | t value | Pr(>\|t\|) |
|---|---|---|---|---|
| **Intercept** | 7.112 | 3.814 | 1.865 | 0.111 |
| **Ploidy** | 3.098 | 1.294 | 2.394 | 0.054 |
| **Years in culture** | -0.290 | 0.098 | -2.956 | 0.025 |

Multiple R-squared: 0.6461; Adjusted R-squared: 0.5282
F-statistic: 5.478 on 2 and 6 DF, p-value: 0.04431

**Table S7**: We used the experimentally measured clone frequencies to calculate a-posteriori saturation curves of scDNA-Seq library sizes for each cell line as previously described (Ruli Gao et al., Nature Genetics 2016). Minimum number of cells required to keep the risk of observing fewer than five cells per clone below 0.01, was calculated as previously described[3], based on a multinomial distribution (3rd column). The number of G0/G1 cells actually sequenced was greater than that minimum for all cell lines, suggesting that all cell lines were sequenced at or above saturation.

| Cell line | Number of clones | Minimum cells | Sequenced G0/G1 cells |
|---|---|---|---|
| SNU-16 | 11 | 279 | 477 |
| KATOIII | 5 | 274 | 604 |
| HGC-27 | 5 | 233 | 576 |
| SNU-668 | 10 | 759 | 1009 |
| NUGC-4 | 4 | 264 | 656 |
| SNU-601 | 12 | 698 | 1186 |
| SNU-638 | 4 | 126 | 537 |
| NCI-N87 | 4 | 451 | 742 |
| MKN-45 | 2 | 149 | 533 |

**Table S8**: Top pathways differentially expressed among the three confirmed epithelial subclones of patient P5931. First three columns list the median pathway activity among members of the three clones, as calculated by GSVA. The p-value of differential activity is shown in the last column (Anova: P<0.003).

| | P5931_Clone6157 | P5931_Clone6155 | P5931_Clone6156 | Anova: P-value |
|---|---|---|---|---|
| Pre-NOTCH Expression and Processing | -0.118 | -0.037 | 0.025 | 1.13E-06 |
| Interleukin-4 and 13 signaling | -0.043 | 0.019 | 0.050 | 2.78E-06 |
| Endosomal Sorting Complex Required For Transport (ESCRT) | 0.034 | -0.029 | -0.100 | 7.88E-06 |
| Signaling by NOTCH | -0.104 | -0.048 | -0.019 | 3.71E-05 |
| Pre-NOTCH Transcription and Translation | -0.121 | -0.046 | 0.025 | 7.22E-05 |
| Synthesis of PIPs at the Golgi membrane | -0.117 | 0.070 | 0.071 | 8.81E-05 |
| Constitutive Signaling by Ligand-Responsive EGFR Cancer Variants | 0.152 | 0.035 | -0.119 | 0.00019 |
| Signaling by EGFR in Cancer | 0.152 | 0.035 | -0.119 | 0.00019 |
| Signaling by Ligand-Responsive EGFR Variants in Cancer | 0.152 | 0.035 | -0.119 | 0.00019 |
| Unfolded Protein Response (UPR) | -0.113 | -0.077 | -0.016 | 0.00038 |
| rRNA processing | 0.425 | 0.328 | -0.263 | 0.00083 |
| rRNA processing in the nucleus and cytosol | 0.437 | 0.334 | -0.275 | 0.00087 |
| Butyrate Response Factor 1 (BRF1) binds and destabilizes mRNA | -0.258 | -0.169 | -0.138 | 0.00089 |
| Major pathway of rRNA processing in the nucleolus and cytosol | 0.454 | 0.348 | -0.292 | 0.00095 |
| EGFR downregulation | 0.065 | -0.067 | -0.106 | 0.00095 |
| Laminin interactions | -0.258 | -0.064 | -0.061 | 0.00119 |
| Influenza Viral RNA Transcription and Replication | 0.557 | 0.500 | -0.275 | 0.00124 |
| Influenza Infection | 0.517 | 0.461 | -0.279 | 0.00131 |
| Influenza Life Cycle | 0.522 | 0.484 | -0.274 | 0.00137 |
| Metabolism | 0.007 | -0.005 | -0.033 | 0.00173 |
| Selenoamino acid metabolism | 0.611 | 0.521 | -0.302 | 0.00201 |
| SRP-dependent cotranslational protein targeting to membrane | 0.584 | 0.507 | -0.284 | 0.00205 |
| E3 ubiquitin ligases ubiquitinate target proteins | -0.031 | -0.011 | -0.092 | 0.00206 |
| Constitutive Signaling by EGFRvIII | 0.035 | -0.100 | -0.179 | 0.00211 |
| Signaling by EGFRvIII in Cancer | 0.035 | -0.100 | -0.179 | 0.00211 |
| Misspliced GSK3beta mutants stabilize beta-catenin | -0.198 | -0.094 | -0.038 | 0.00240 |
| S33 mutants of beta-catenin aren't phosphorylated | -0.198 | -0.094 | -0.038 | 0.00240 |
| S37 mutants of beta-catenin aren't phosphorylated | -0.198 | -0.094 | -0.038 | 0.00240 |
| S45 mutants of beta-catenin aren't phosphorylated | -0.198 | -0.094 | -0.038 | 0.00240 |
| T41 mutants of beta-catenin aren't phosphorylated | -0.198 | -0.094 | -0.038 | 0.00240 |
| phosphorylation site mutants of CTNNB1 are not targeted to the proteasome by the destruction complex | -0.198 | -0.094 | -0.038 | 0.00240 |
| Nonsense Mediated Decay (NMD) enhanced by the Exon Junction Complex (EJC) | 0.588 | 0.513 | -0.322 | 0.00241 |
| Nonsense-Mediated Decay (NMD) | 0.588 | 0.513 | -0.322 | 0.00241 |
| Depolymerisation of the Nuclear Lamina | -0.163 | -0.045 | 0.000 | 0.00277 |
| Initiation of Nuclear Envelope Reformation | -0.236 | -0.082 | -0.024 | 0.00288 |
| Nuclear Envelope Reassembly | -0.236 | -0.082 | -0.024 | 0.00288 |
| Nonsense Mediated Decay (NMD) independent of the Exon Junction Complex (EJC) | 0.658 | 0.594 | -0.242 | 0.00293 |
